# Adaptation to heat and ocean fertilization, two keys for understanding the massive *Sargassum* growth in the Atlantic

**DOI:** 10.64898/2025.12.08.692933

**Authors:** Roberto Velázquez-Ochoa, Susana Enríquez

**Affiliations:** Laboratory of Photobiology and Functional Allometry, Unidad Académica de Sistemas Arrecifales Puerto Morelos, Instituto de Ciencias del Mar y Limnología, Universidad Nacional Autónoma de México, Apdo. Postal 13, Cancun, Mexico

**Keywords:** Algal blooms, Brown algae, Comparative ecophysiology, Functional traits, Global warming, Photosynthesis, Optical traits

## Abstract

- A floating ecosystem constituted by three genetic variants of holopelagic *Sargassum* has extended since 2011 throughout the tropical North Atlantic without spatial restrictions.
- We characterized the differential capacity and efficiency of each variant to collect light and fix this energy in photosynthesis under variable light and temperature regimes, focusing on the description of key physiological and optical traits the differential response to light and temperature.
- Our results revealed metabolic adaptations of two genetic variants to the warmer conditions of the tropical Atlantic and contrasting efficiencies in light absorption and use in photosynthesis, indicative of distinct competitive abilities under growth limitations.
- We concluded that the increased fertility of a warmer ocean is the most plausible explanation for the massive presence of holopelagic *Sargassum* in the tropical Atlantic, which also may explain the current ecological success of the opportunistic strategy of a previously rare variant. The optical and physiological descriptors documented can assist in developing quantitative models for predicting *Sargassum* biomass in the Atlantic.

## Introduction

In the middle of the Atlantic gyre (20°–40°N and 30°–75°W), holopelagic *Sargassum* has formed an emblematic and floating oceanic ecosystem, the Sargasso Sea (Winge 1923). The gyre keeps recirculating the floating algal biomass, restricting its expansion in the Atlantic Ocean and providing habitat, shelter and food for a high biodiversity of species (Huffard et al. 2014). Initially, eight different drifting *Sargassum* species/variants were described in the North Atlantic (Taylor 1960), although only two species, *S. fluitans* and *S. natans,* are currently accepted (Taylor 1960; Stiger-Pouvreau et al. 2023), with *S. natans* I and *S. fluitans* III being the most common variants (Parr 1939). Since 2011, another floating *Sargassum* population has been reported for the tropical Atlantic around the North Equatorial Recirculation Region (NERR) (Wang et al. 2019; Jouanno et al. 2023). This floating ecosystem, named the “*Great Atlantic Sargassum Belt*” (GASB), which extends as a belt along the Atlantic Ocean reaching extensions up to ∼5.5 million km^2^, is formed by the two common variants in the Sargasso Sea and a rare type, *S. natans* VIII (Gower and King 2011; Schell et al. 2015; Amaral Zettler et al. 2016). Fresh weight maxima of ∼20 million tons (Wang et al. 2019) and >22 million tons (Jouanno et al. 2023) have been reported for the GASB in, respectively, July 2018 and July 2021. Notably, a new historical record of 38 milllion tons was reached in July 2025 (https://optics.marine.usf.edu/projects/SaWS/pdf/Sargassum_outlook_2025_bulletin07_USF.pdf). For comparison and despite the large spatial, seasonal and interannual variability in *Sargassum* biomass (Gower and King 2011; Niermann 1986) and productivity (Lapointe 1995) documented for the Sargasso Sea, the estimated maximum production (Winge 1923; Lomas et al. 2013) is less than 10 million tons over an area of ∼4.16 million km^2^, for an algal community that survives under strong nutrient limitations (Lapointe 1995; Cavender-Bares et al. 2001) and within a temperature range between 19°–29°C (Lomas et al. 2013).

The origin and establishment of such a productive population of holopelagic *Sargassum* in the NERR is still under discussion (Wang et al. 2019; Oviatt et al. 2019; Johns et al. 2020; Jouanno et al. 2025; Jung et al. 2025), as well as its recirculation patterns through different oceanic routes (Putman et al. 2018) or the large spatial and temporal variability in variant dominance (Schell et al. 2015; Amaral Zettler et al. 2016). As nutrients stimulate *Sargassum* growth and photosynthesis (Lapointe 1995), it has been proposed that oceanic fertilization from western (i.e., Amazon River) and/or eastern (i.e., Congo River) Atlantic regions may explain *Sargassum* proliferation in the NERR (Wang et al. 2019; Oviatt et al. 2019), or equatorial upwelling of phosphorus (Jung et al. 2025). Other hypotheses for GASB formation have suggested the dispersion of a substantial floating biomass from the Sargasso Sea towards lower latitudes after a climatological anomaly in winter 2009–2010 (Johns et al. 2020; Jouanno et al. 2025), or due to the direct effects of climate change. Optimal conditions in the NERR for *S. natans* VIII could also explain the proliferation of this variant in the tropical Atlantic, but this, together with why *S. fluitans* III is also so successful in the NERR, remains poorly understood.

The genus *Sargassum* (Phaeophyta, order *Fucales*) displays a large diversity of species and complex morphologies with a cosmopolitan distribution across tropical, subtropical and temperate marine environments, where they exhibit the highest diversity and abundance (Yip et al. 2020; Stiger-Pouvreau et al. 2023). Some species are highly productive with elevated competitive abilities for light, nutrients and space (Stiger-Pouvreau et al. 2023), large phenotypic plasticity and resilience to environmental stress, and the ability to disperse over long distances through propagule drifting with diverse reproductive patterns (Doust and Doust 1988; Stiger-Pouvreau et al. 2023). These features give them the potential to become aggressive invaders and cause blooms and severe ecological disorders (Salvaterra et al. 2013; Stiger-Pouvreau et al. 2023). However, most of the *Sargassum* species are benthic, distributed across a broad depth range. Only two species have colonized the ocean surface, completing their life cycle unattached to a substrate and exposed to air and full light (Jerzmańska et al. 1976; Yip et al. 2020; Stiger-Pouvreau et al. 2023). Offshore pelagic algae in the central ocean are adapted to survive under very poor nutrient conditions (Lapointe 1995). These populations share similar genetic lineages with benthic species (González-Nieto et al. 2020; Jouanno et al. 2025), although they lack sexual reproduction (Stiger-Pouvreau et al. 2023), a character that strongly affects their adaptive abilities. It is worth to note that despite these adaptive limitations, consistent morphological (Siuda et al. 2024) and genetic (Amaral Zettler et al. 2016; González-Nieto et al. 2020; Siuda et al. 2024) variability has been documented for the three holopelagic variants that form the GASB, with no intermediate form (Amaral Zettler et al. 2016; Siuda et al. 2024). Differences in blade and vesicle size and shape allow distinction among them (Siuda et al. 2024), whereas minor but also consistent genetic variability supports their genetic divergence, greater for *S. natans* and *S. fluitans* and smaller for the two *natans* types (Amaral Zettler et al. 2016;). Differences in relative growth rates (RGR, day^-1^) have also been reported (Hanisak and Samuel 1987; Corbin and Oxenford 2023), associated with their contrasting responses to nutrients (Lapointe 1986; 1995), optimal temperature (Hanisak and Samuel 1987), and salinity tolerance (Corbin and Oxenford 2023). It has been proposed separating these three variants into different taxonomic entities or species (Amaral Zettler et al. 2016), but the maintenance of the ‘variety’ classification has been also recommended (Siuda et al. 2024) until the speciation stage of these taxa can be better understood, as current analyses are restricted to mitochondrial and plastid markers. Understanding the variability of nuclear genomes will help elucidate how morphological differences have been fixed through asexual lineages.

We characterized here the optical and photo-physiological/metabolic traits of *S. natans* I, *S. natans* VIII, and *S. fluitans* III, following previous morpho-functional approaches (Enríquez and Sand-Jensen 2003; Cayabyab and Enríquez 2007; Schubert et al. 2015; Vásquez-Elizondo and Enríquez 2016; 2017). Our aim was to contribute to the comprehension of these genetic differences by identifying key variability in their capacity and efficiency to collect light and fix this energy as organic carbon through photosynthesis, under different light and temperature regimes. Knowing contrasting functional differences between variants will improve our understanding of the spatial and temporal variability of the GASB, deepening in the comprehension of why this population of floating algae was established at lower latitudes and higher temperatures. This information may also facilitate the development of quantitative/predictive models of *Sargassum* biomass from light and temperature satellite data (Behrenfeld and Falkowski 1997). To scale the experimental results to a broader context, three comparative analyses were also performed for the variation in the relative abundance, algal size, and growth rates (RGR, day^-1^) of the three *Sargassum* variants, compiling old and more recent literature data (Supporting Information tables S1-S3).

## Materials and Methods

### Sample collection

Fresh samples of *Sargassum natans* I and VIII and *Sargassum fluitans III (*hereafter referred to only as *S. fluitans*) were collected during the 2017–2020 summers in a Mexican Caribbean reef lagoon, Puerto Morelos (20° 52’ N, 86° 50’ W), before algal arrival on the beach. Species and morphotype identifications were based on structural features following the descriptions of Schell et al (2015) and Amaral-Zettler et al (2016). The samples were placed in the UNAM-UASA mesocosm system and fed directly with seawater from the reef lagoon (filtered through 0.45-pm filters). The annual average temperature of the lagoon water was previously documented by Scheufen et al. (2017) to be at 28°C. The selected samples for experimental analysis were cleaned of epiphytes, epibionts and sand and placed in 150 L tanks with two underwater pumps that allowed maintenance of constant water circulation. The irradiance was shaded with neutral meshes and maintained at 80% of the surface irradiance (E_s_). Experimental analyses were performed no more than three days after sample collection to prevent the induction of new photoacclimatory conditions in the organisms. Thallus determinations were performed on independent phylloides (i.e., algal structures, blades, that look like leaves).

### Morphological characterization

Fifty fresh phylloides (blades) without any signs of herbivory or damage were randomly selected and scanned to measure blade size via digital tools (IMAGE−J software). No differences were detected in terms of blade size between different branch orders, as documented in benthic species (Abbott 1992). Thallus size was expressed as area (cm^2^), or dry weight (mg DW), which allowed calculation of a common plant trait, specific leaf area (SLA, cm^2^ g DW^−1^). For dry weight determination, fragments were placed in a drying oven for 24 hours at 80°C until a constant weight was reached, after which they were weighed with an analytical balance. We maintained the same notation, SLA, for a known plant trait, rather than generating a new notation and name (specific thallus area) for the same descriptor.

The pigment content was determined spectrophotometrically using a miniature Ocean Optics USB4000 (Ocean Optics Ltd., FL) on the same samples used for light absorption determination and for the same thallus fragments used for photosynthetic measurements. These samples were immediately frozen (−70°C) after measurement until a set of samples was formed for pigment analysis. Aqueous acetone 90% (vol:vol) was used for the extraction of chlorophyll *a* (Chl*a*) and chlorophyll *c* (Chl*c*) after the tissue was ground in a mortar using a manual pistile under cold, low-light conditions. The samples were homogenized in darkness at 4°C for 12 hours and then centrifuged for 2 minutes at 130,000 RPM in a 3 ml blank cuvette. The equations of Jeffrey and Humphrey (1975) for brown algae were used to calculate the chlorophyll contents. For each sample, the dry weight and projected surface area were determined and used to normalize the pigment content to the area and weight.

### Optical determinations

Determinations of light transmission (absorbance, D) were performed according to Vásquez-Elizondo et al. (2017) using a miniature Ocean Optics USB2000 spectroradiometer (Ocean Optics, Inc., Dunedin, FL, USA) equipped with an opal glass in front of the detector (Shibata 1959). The blades (phyllodes) were fragmented to exclude the midribs, which are thicker than the rest of the blade and contribute to increase residual scattering and light loss during optical determination. Light absorption spectra were performed between 400 and 750 nm at intervals of 1 nm using OceanView software. The light absorption capacity of the thallus or absorptance (i.e., fraction of incident light absorbed) was estimated by subtracting the reflectance (R, fig S1 in Supplementary Materials) from absorbance (D) determinations according to Vásquez-Elizondo et al. (2019), via the following equation:

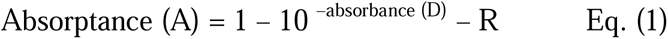

Absorptance was calculated for the PAR (400–700 nm) average (A_PAR_) and for the chlorophyll *a* (Chl*a*) peak at 675 nm (A_Chl*a*_). The variation in the efficiency of pigments for absorbing light was assessed by estimating the specific absorption coefficients, for Chl*a* (a*_Chl*a*_; m^2^ mg Chl*a*^−1^) and for the PAR average (a*_PAR_; m^2^ mg Chl[*c* + *a*]^−1^). These estimations were performed via the following equation:

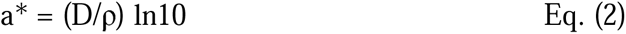

where D represents the absorbance value (D_PAR_ and D_Chl*a*_) and ρ represents the total pigment cross section in mg m^-2^ (Chl*a* for a*_Chl*a*_, and total chlorophyll content Chl[*c* + *a*] for a*_PAR_).

### Physiological determinations

#### A Response to light

Oxygen sensors (OXSP5, PyroScience) were used for the determinations of the photosynthesis vs. irradiance (P–E) curves (n = 5 and 10 replicates per curve). The sensors were connected to a closed custom acrylic water chamber (670 ml) with filtered seawater (0.45 µm) and a data logger (FireSting-O_2_) that was operated with Pyro Oxygen Logger software. We calibrated the oxygen sensor spot twice, first with 21% air saturation and then bubbling the water with N_2_ gas to approach 0% O_2_ concentration. We did this by following the steps outlined in the Oxygen Sensor Fiber-optic and Contactless, Document Version 1.015 by PyroScience (https://www.manualslib.com/manual/2284456/Pyroscience-Pico-T.html). Samples were located in 706 mL home-made cylindrical acrylic chambers (10 cm of diameter and 9 cm heigh, see fig. S2), developed originally for photosynthesis determinations on corals. Several phylloides (see fig. S2A) were placed on a plastic net, 5 cm of diameter, framed as a 7 cm heigh PVC table, fixed to the chamber. This allowed maintaining the samples at the same distance to the light source, in a fixed position, preventing phylloide overlapping (fig. S2B). The plastic net where the samples were placed, provides an open surface to facilitate water circulation around the samples during the incubations. Moderate agitation was generated and controlled individually in each chamber, by a magnetic stirring system. All the chambers were enclosed by a water jacked, connected to a recirculating bath with a controlled temperature system (Thermo Fisher Scientific, Boston, USA). The number of blades used in each incubation changed between 9-16 for the smallest variant, *S. natans* I, to 4-7 for the largest, *S. natans* VIII (fig. S2C). The selected thalli were first dark acclimated for 15 minutes, and then progressively exposed to 13 different light intensities of 8, 14, 21, 37, 61, 104, 122, 238, 344, 593, 866, 1123, and 2245 µmol photons m^2^ s^−1^. The exposure time for each intensity was ≈5 minutes. After the light incubations, the samples were newly exposed to darkness to measure ‘light-enhanced respiration’ or post-illumination respiration (R_L_), generally larger than the respiration of dark acclimated samples (R_D_). Photosynthetic parameters derived from the P–E curves were calculated following Cayabyab and Enríquez (2007). Dark respiration (R_D_, µmol O_2_ cm^−2^ h^−1^) was estimated from the intercept of the regression line on the ordinate; photosynthetic efficiency (α, µmol O_2_ cm^−2^ h^−1^ / µmol incident quanta^−1^ m^−2^ s^−1^) was estimated from the initial slope via linear least-squares regression analysis; the maximum net photosynthetic rates (NP_max_, µmol O_2_ cm^−2^ h^−1^) was determined from the average values of measurements under saturation irradiance; and gross photosynthesis, GP_max_, was calculated by adding the absolute R_L_ values to net P_max_. The saturation irradiance (E_k_, µmol quanta m^−2^ s^−1^) was calculated as the ratio P_max_/α; and the compensation irradiance (E_c_, µmol quanta m^−2^ s^−1^) was calculated as the intercept with the irradiance axis of α, the initial slope at sub-saturating light. The photosynthesis: respiration ratios (P:R) were estimated by dividing the maximum gross photosynthetic rates (GP_max_, µmol O_2_ cm^−2^ h^−1^) by the light-enhanced respiration rate (R_L_, µmol O_2_ cm^−2^ h^−1^).

All photosynthetic determinations were normalized to area (µmol O_2_ cm^−2^ h^−1^) and dry weight (µmol O_2_ g DW h^−1^). The maximum quantum yield of photosynthesis (Φ*_max_,* mol O_2_ molecules evolved per mol quanta absorbed) was also estimated for the same thalli used in the optical determinations, after absorptance correction of the photosynthetic efficiency, α, as α/A (Cayabyab and Enríquez 2007; González-Guerrero et al. (2021). The minimum quantum requirements of thallus photosynthesis were calculated as the inverse of the quantum yield (*1/*Φ*_max_*; mol quanta absorbed per mol O_2_ evolved).

#### B Response to temperature

P–E curves were generated for seven distinct temperatures, ranging from 22°C to 34°C with 2°C intervals. The temperature range selected reflects the annual variation observed in the reef lagoon of Puerto Morelos, located in the northern region of the Mexican Caribbean, with the addition of +4°C to examine the response of *Sargassum* to temperatures above the maximum average value of 30°C in summer (Scheufen et al. 2017). The samples were pre-incubated before the physiological determinations. For 60 minutes, for the temperatures between 24-30°C; and for 120 minutes, for the temperatures outside of the optimal range such as 22°, 32°, and 34°C. The number of replicates for each temperature was between 5 and 10 determinations, depending on the variability found among samples. We additionally used 3–9 more replicates for R_D_, NP_max_, and R_L_ determinations once E_K_ was known. To determine R_D_, these samples were exposed first to darkness for 15 minutes and then to saturating irradiance levels (above E_K_) for 30 minutes to determine NP_max_, followed by a final exposure to darkness to determine R_L_. For the calculation of the metabolic scaling factor, Q_10_, we used linear regression models according to (Vásquez-Elizondo and Enríquez 2016; Scheufen et al. 2017), estimated as the slope of the linear increase between 24°C and 34°C (10 degrees).

### Data analysis

Most results are expressed as the mean ± standard error (SE). One-way ANOVA was used to compare optical and morphological traits. Tukey post hoc tests enabled the identification of significant differences among species and/or treatments. Analyses of covariance (ANCOVA-tests) were also used to test for significant differences either in the intercept or the slope of the linear least squares regression associations. The data were analysed using R-software (R Core Team 1964) and Kaleidagraph software.

## Results

### Morphological and optical characterizations

The morphological description confirmed that *S. natans* VIII is the variant with the largest blades and *S. natans* I is the smallest (**Fig. 1A**). The intermediate size of *S. fluitans* III (hereafter referred to only as *S. fluitans*) also presented the largest ‘biomass extension’ (SLA= 165 ± 3.58 cm^2^ gDW^-1^; **Fig. 1B**; **Table 1**). Thallus pigmentation did not vary significantly among the variants (**Fig. 1C**), and light absorption spectra exhibited the representative shape of Phaeophyta (**Fig. 2**). Despite these similarities, the average PAR absorptance (fraction or % of incident light absorbed by photosynthetic pigments, A_PAR_) was greatest for the blades of *S. natans* I, the variant with the largest variability in pigmentation (**Fig. 3A**). These differences were also observed for the Chl*a* absorption peak at 675 nm (A_Chl*a*_; **Table 1**). *S. natans* VIII, differed from *S. fluitans* in the Chl[*c/a*] molar ratio (**Table 1**), and presented the smallest peak at 675 nm and the highest light absorption around 405-414 nm, with a peak at approximately 412 nm. This highlights significant differences for *S. natans* VIII in the proportion of blue to red light absorbed (**Fig. 2**).

**Figure 1:**
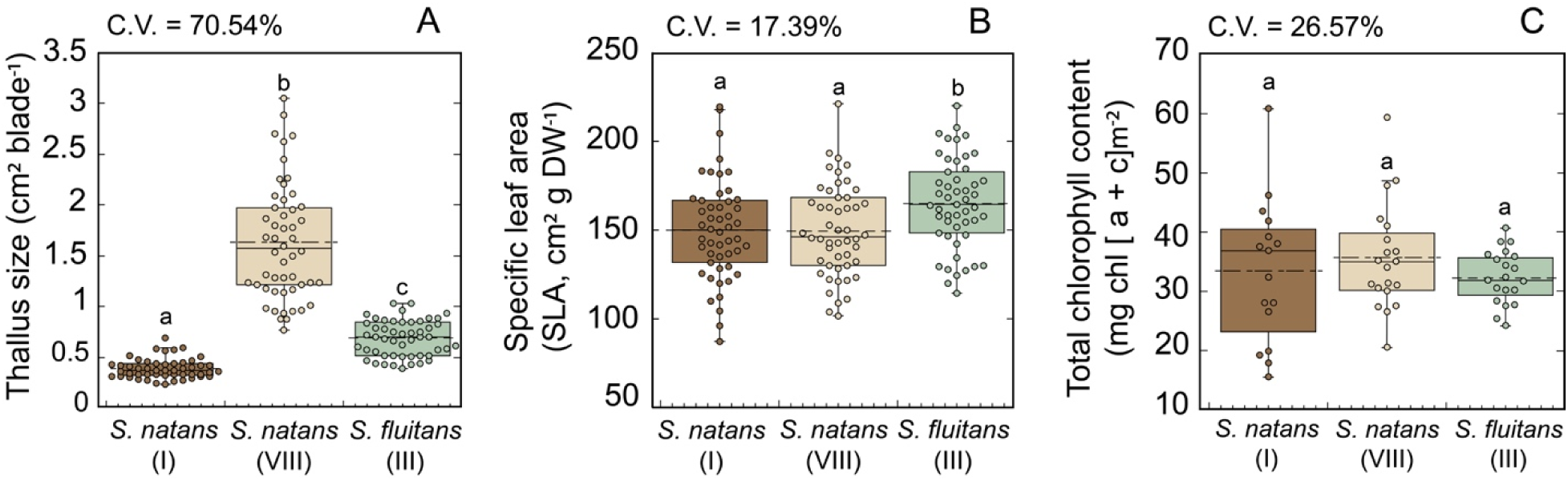
Variability of the structural traits characterized for the three genetic variants of holopelagic *Sargassum* investigated. Boxplots displaying the structural variability in (A) thallus size (cm^2^ blade^-1^); (B) thallus specific area, SLA (cm^2^ g DW^−1^); and (C) thallus pigmentation (chlorophyll content per unit of area; mg Chl [a+c] m^−2^).

**Figure 2.**
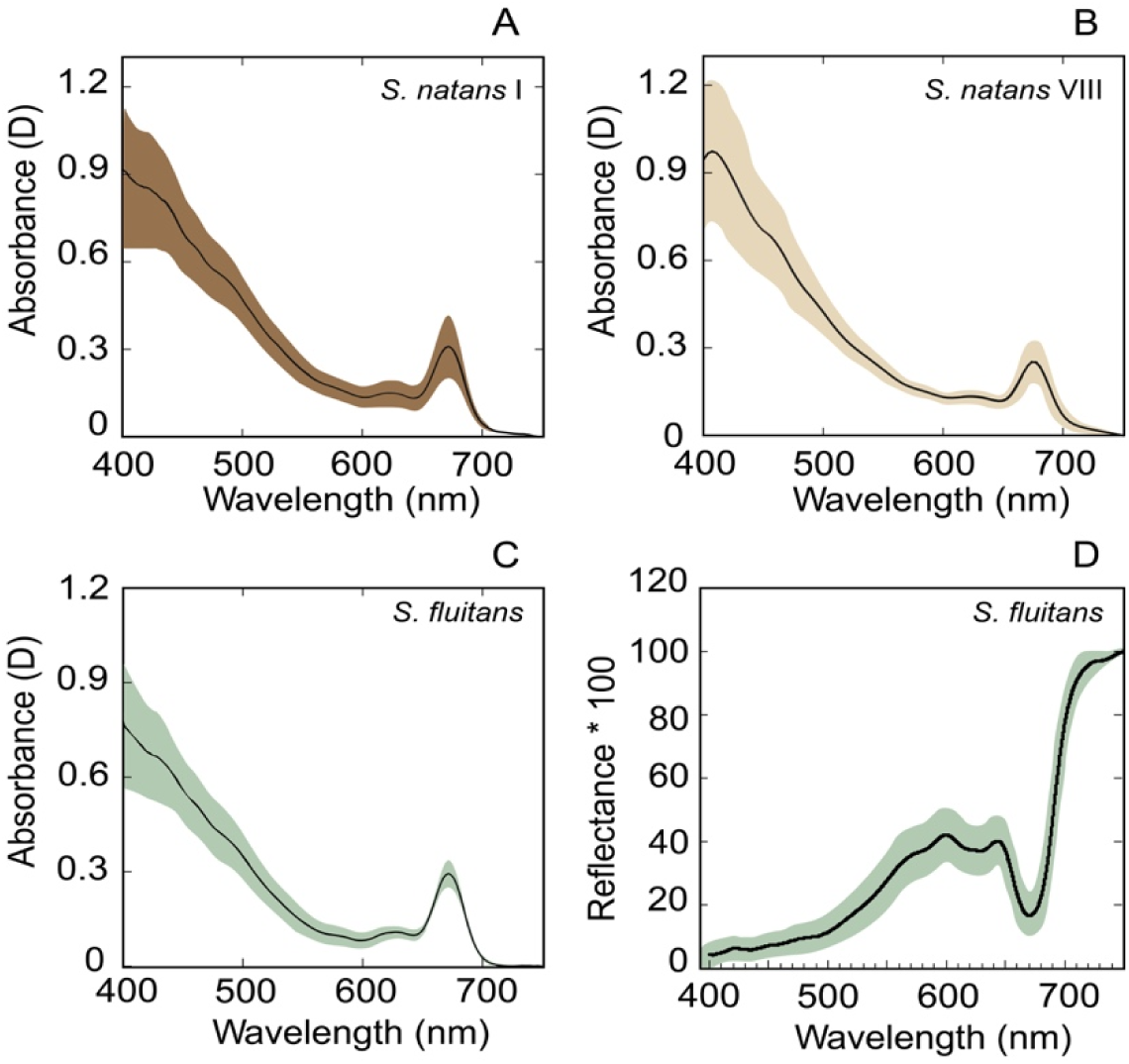
Light absorption and reflection spectra for holopelagic *Sargassum*. Plots A–C illustrate the light absorption spectra of *S. natans* I (A); *S. natans* VIII (B) and *S. fluitans* (C). Plot D describes the reflectance spectrum of *S. fluitans*. No reflectance differences were observed among variants. The solid line represents the average spectrum of independent replicated spectra (n= 9-16), and the shaded area represents the standard deviation variability.

**Figure 3.**
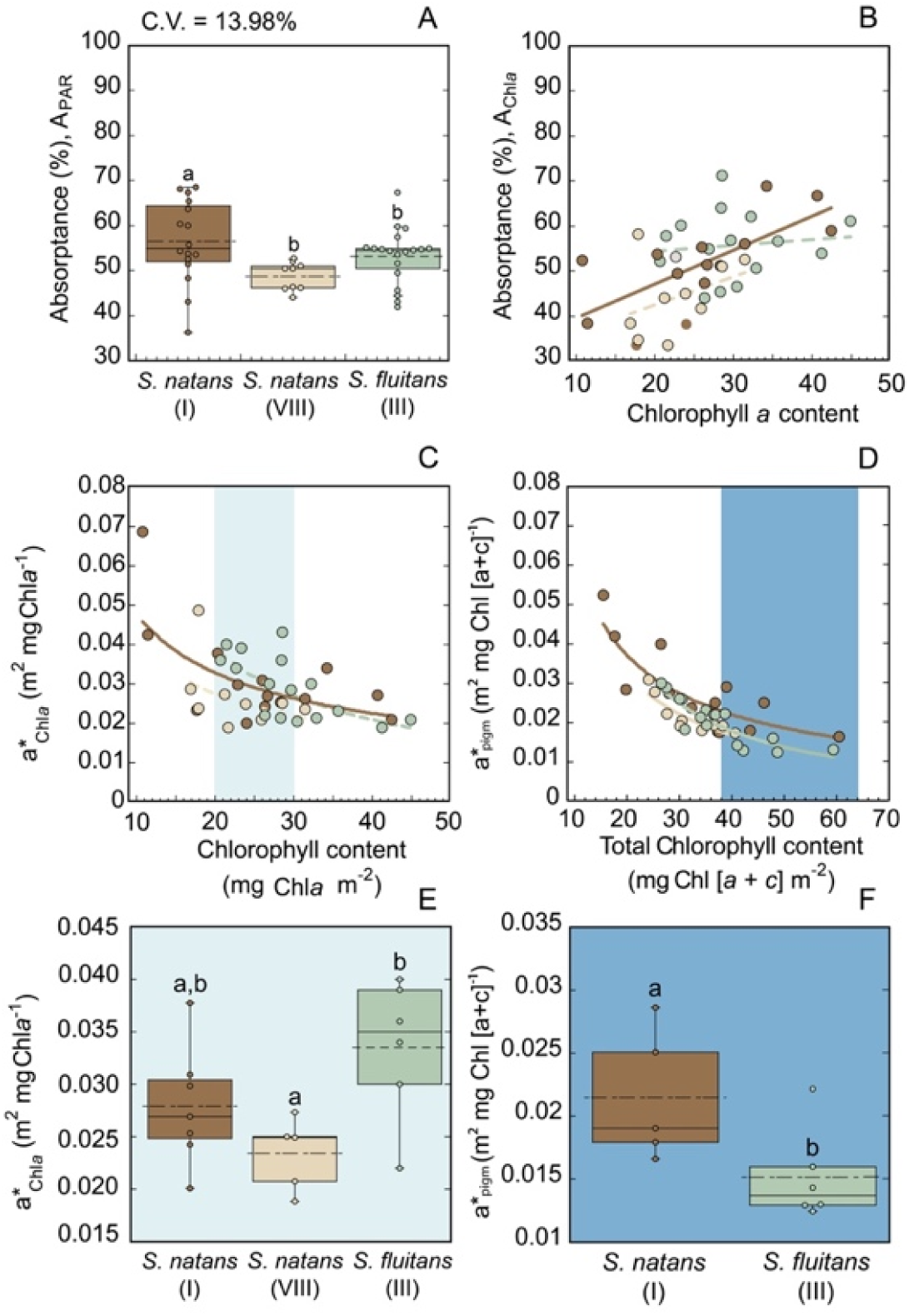
Variability of the structural and optical traits of the three genetic variants of holopelagic *Sargassum* investigated. (A) Boxplot displaying the variability in absorptance for the PAR average (A_PAR_). (B) Variability of A_PAR_ as a function of the variation in thallus pigmentation; and (C) and (D) describe, respectively, the variability of a*_Chl*a*_ against Chl*a* content; and a*_PAR_ against total Chl [a+c] content. Boxplots in (E) and (F) describe, respectively, the differences in a*_Chl*a*_ and in a*_PAR_, for thalli with intermediate Chl*a* values (in light blue); and for the variants of *S. natans* I and *S. fluitans,* with the largest pigmentation (> 40 mg Chl [a+c] m^−2^; in dark blue). Different letters indicate significant differences at the < 0.05 level (Tukey Post-Hoc test). Box plots encompass the 25 and 75% quartiles, whereas the central solid line corresponds to the median, and the discontinued line represents the mean. In plots B-D dots indicate the value of each replicate characterized, solid lines represent significant association of variation for linear (B) and power functions (C, D) (p<0.05). Colors are maintained consistent for each genetic variant, dark brown for *S. natans* I, light brown for *S. natans* VIII and green for *S. fluitans* III.

**Table 1:**
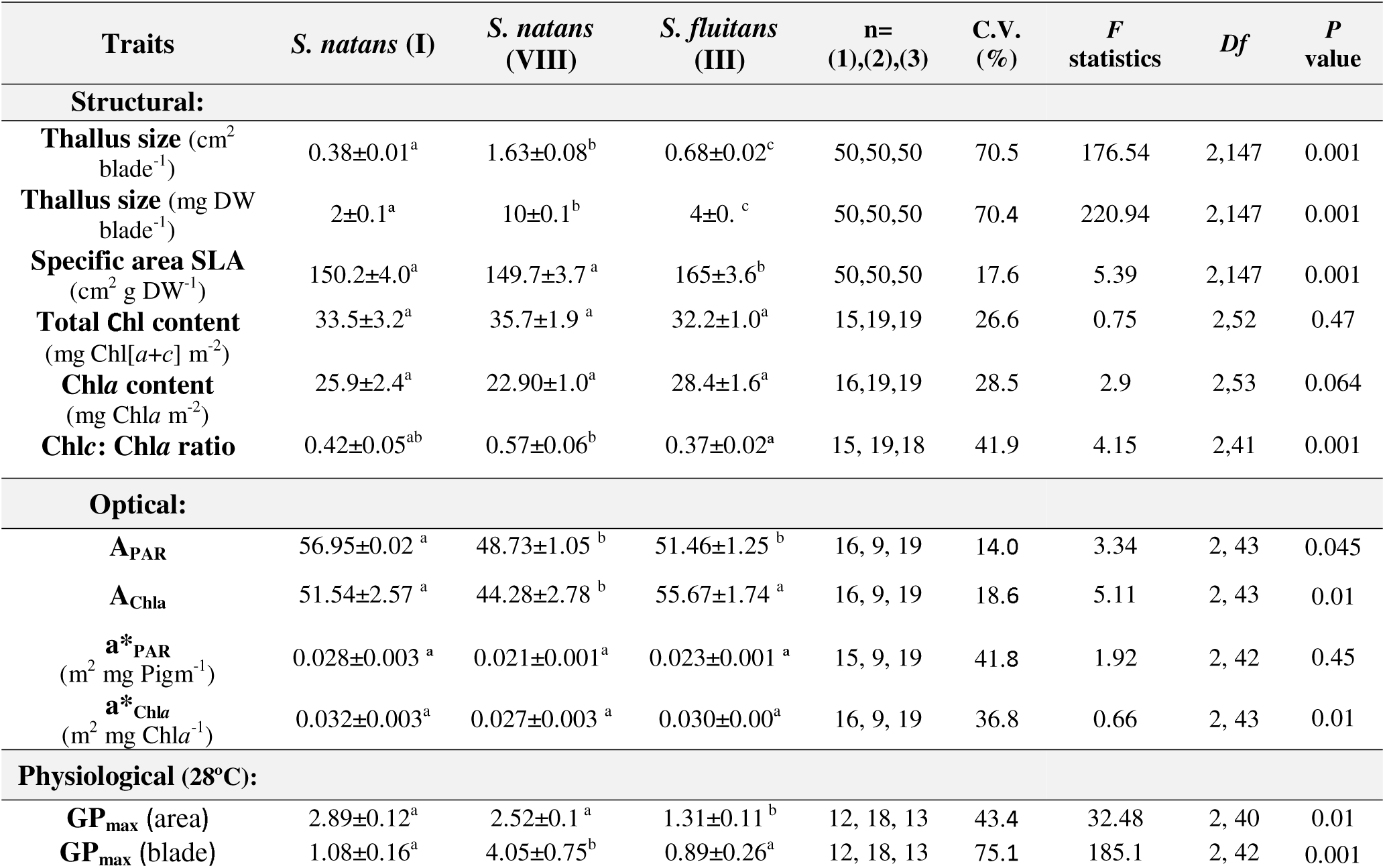
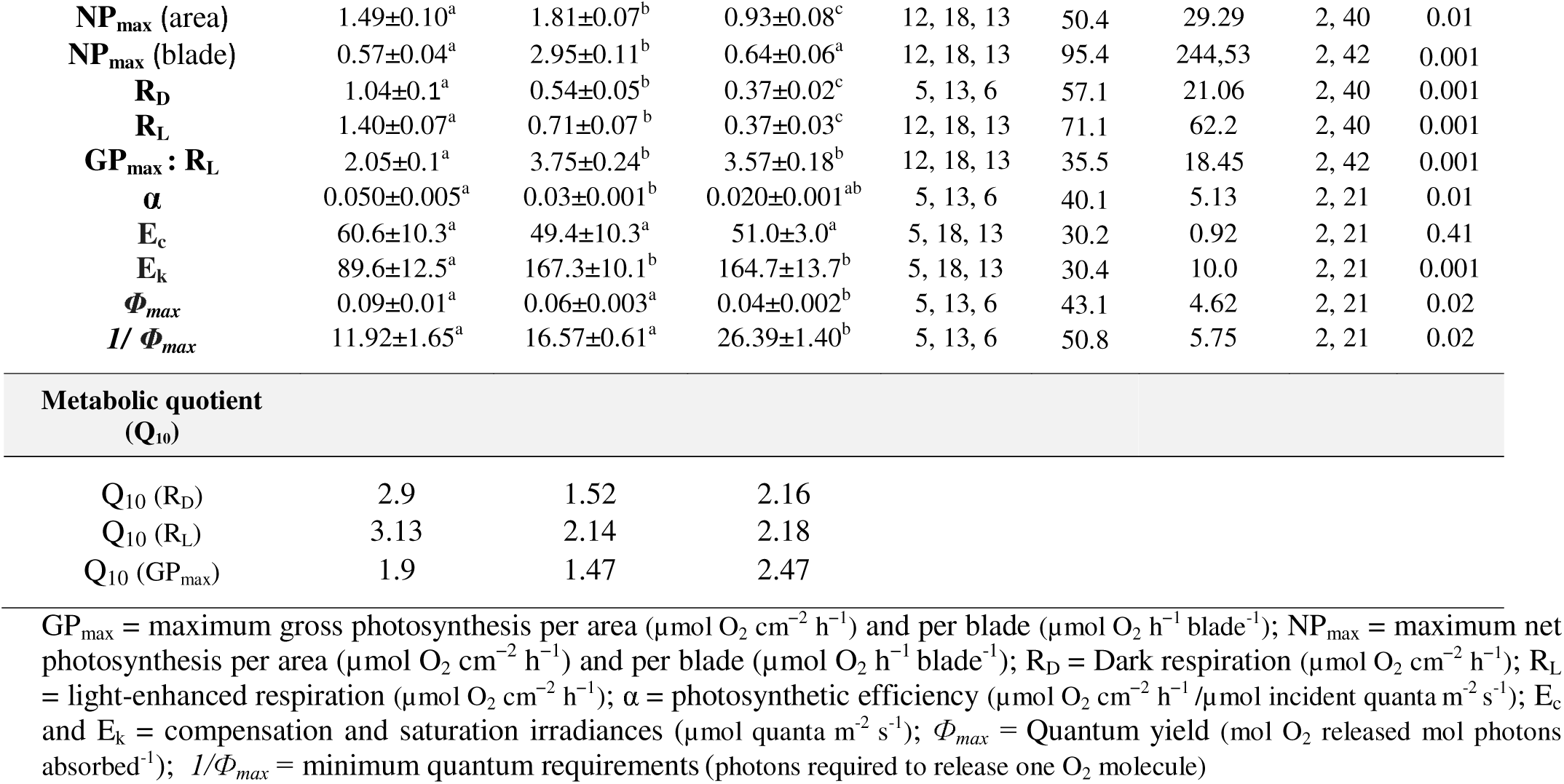
Mean ± SE of the structural, optical and physiological traits characterized in this study for the three genetic variants of holopelagic *Sargassum* collected in the Mexican Caribbean. Different letters indicate significant differences at the 0.05 level (Tukey Post-Hoc test).

| Traits | *S. natans* (I) | *S. natans* (VIII) | *S. fluitans* (III) | n=(1),(2),(3) | C.V. (%) | F statistics | Df | P value |
| --- | --- | --- | --- | --- | --- | --- | --- | --- |
| **Structural:** |  |  |  |  |  |  |  |  |
| **Thallus size** (cm<sup>2</sup> blade<sup>-1</sup>) | 0.38±0.01<sup>a</sup> | 1.63±0.08<sup>b</sup> | 0.68±0.02<sup>c</sup> | 50,50,50 | 70.5 | 176.54 | 2,147 | 0.001 |
| **Thallus size** (mg DW blade<sup>-1</sup>) | 2±0.1<sup>a</sup> | 10±0.1<sup>b</sup> | 4±0.1<sup>c</sup> | 50,50,50 | 70.4 | 220.94 | 2,147 | 0.001 |
| **Specific area SLA** (cm<sup>2</sup> g DW<sup>-1</sup>) | 150.2±4.0<sup>a</sup> | 149.7±3.7<sup>a</sup> | 165±3.6<sup>b</sup> | 50,50,50 | 17.6 | 5.39 | 2,147 | 0.001 |
| **Total Chl content** (mg Chl[a+c] m<sup>-2</sup>) | 33.5±3.2<sup>a</sup> | 35.7±1.9<sup>a</sup> | 32.2±1.0<sup>a</sup> | 15,19,19 | 26.6 | 0.75 | 2,52 | 0.47 |
| **Chla content** (mg Chla m<sup>-2</sup>) | 25.9±2.4<sup>a</sup> | 22.90±1.0<sup>a</sup> | 28.4±1.6<sup>a</sup> | 16,19,19 | 28.5 | 2.9 | 2,53 | 0.064 |
| **Chlc: Chla ratio** | 0.42±0.05<sup>ab</sup> | 0.57±0.06<sup>b</sup> | 0.37±0.02<sup>a</sup> | 15, 19,18 | 41.9 | 4.15 | 2,41 | 0.001 |
| **Optical:** |  |  |  |  |  |  |  |  |
| **A<sub>PAR</sub>** | 56.95±0.02<sup>a</sup> | 48.73±1.05<sup>b</sup> | 51.46±1.25<sup>b</sup> | 16, 9, 19 | 14.0 | 3.34 | 2, 43 | 0.045 |
| **A<sub>Chla</sub>** | 51.54±2.57<sup>a</sup> | 44.28±2.78<sup>b</sup> | 55.67±1.74<sup>a</sup> | 16, 9, 19 | 18.6 | 5.11 | 2, 43 | 0.01 |
| **a\*<sub>PAR</sub>** (m<sup>2</sup> mg Pigm<sup>-1</sup>) | 0.028±0.003<sup>a</sup> | 0.021±0.001<sup>a</sup> | 0.023±0.001<sup>a</sup> | 15, 9, 19 | 41.8 | 1.92 | 2, 42 | 0.45 |
| **a\*<sub>Chla</sub>** (m<sup>2</sup> mg Chla<sup>-1</sup>) | 0.032±0.003<sup>a</sup> | 0.027±0.003<sup>a</sup> | 0.030±0.00<sup>a</sup> | 16, 9, 19 | 36.8 | 0.66 | 2, 43 | 0.01 |
| **Physiological (28°C):** |  |  |  |  |  |  |  |  |
| **GP<sub>max</sub>** (area) | 2.89±0.12<sup>a</sup> | 2.52±0.1<sup>a</sup> | 1.31±0.11<sup>b</sup> | 12, 18, 13 | 43.4 | 32.48 | 2, 40 | 0.01 |
| **GP<sub>max</sub>** (blade) | 1.08±0.16<sup>a</sup> | 4.05±0.75<sup>b</sup> | 0.89±0.26<sup>a</sup> | 12, 18, 13 | 75.1 | 185.1 | 2, 42 | 0.001 |
| **NP<sub>max</sub>** (area) | 1.49±0.10<sup>a</sup> | 1.81±0.07<sup>b</sup> | 0.93±0.08<sup>c</sup> | 12, 18, 13 | 50.4 | 29.29 | 2, 40 | 0.01 |
| **NP<sub>max</sub>** (blade) | 0.57±0.04<sup>a</sup> | 2.95±0.11<sup>b</sup> | 0.64±0.06<sup>a</sup> | 12, 18, 13 | 95.4 | 244.53 | 2, 42 | 0.001 |
| **R<sub>D</sub>** | 1.04±0.1<sup>a</sup> | 0.54±0.05<sup>b</sup> | 0.37±0.02<sup>c</sup> | 5, 13, 6 | 57.1 | 21.06 | 2, 40 | 0.001 |
| **R<sub>L</sub>** | 1.40±0.07<sup>a</sup> | 0.71±0.07<sup>b</sup> | 0.37±0.03<sup>c</sup> | 12, 18, 13 | 71.1 | 62.2 | 2, 40 | 0.001 |
| **GP<sub>max</sub> : R<sub>L</sub>** | 2.05±0.1<sup>a</sup> | 3.75±0.24<sup>b</sup> | 3.57±0.18<sup>b</sup> | 12, 18, 13 | 35.5 | 18.45 | 2, 42 | 0.001 |
| **α** | 0.050±0.005<sup>a</sup> | 0.03±0.001<sup>b</sup> | 0.020±0.001<sup>ab</sup> | 5, 13, 6 | 40.1 | 5.13 | 2, 21 | 0.01 |
| **E<sub>c</sub>** | 60.6±10.3<sup>a</sup> | 49.4±10.3<sup>a</sup> | 51.0±3.0<sup>a</sup> | 5, 18, 13 | 30.2 | 0.92 | 2, 21 | 0.41 |
| **E<sub>k</sub>** | 89.6±12.5<sup>a</sup> | 167.3±10.1<sup>b</sup> | 164.7±13.7<sup>b</sup> | 5, 18, 13 | 30.4 | 10.0 | 2, 21 | 0.001 |
| **Φ<sub>max</sub>** | 0.09±0.01<sup>a</sup> | 0.06±0.003<sup>a</sup> | 0.04±0.002<sup>b</sup> | 5, 13, 6 | 43.1 | 4.62 | 2, 21 | 0.02 |
| **I/ Φ<sub>max</sub>** | 11.92±1.65<sup>a</sup> | 16.57±0.61<sup>a</sup> | 26.39±1.40<sup>b</sup> | 5, 13, 6 | 50.8 | 5.75 | 2, 21 | 0.02 |

Metabolic quotient (Q<sub>10</sub>)
|  |  |  |  |
| --- | --- | --- | --- |
| Q<sub>10</sub> (R<sub>D</sub>) | 2.9 | 1.52 | 2.16 |
| Q<sub>10</sub> (R<sub>L</sub>) | 3.13 | 2.14 | 2.18 |
| Q<sub>10</sub> (GP<sub>max</sub>) | 1.9 | 1.47 | 2.47 |

GP<sub>max</sub> = maximum gross photosynthesis per area (μmol O<sub>2</sub> cm<sup>-2</sup> h<sup>-1</sup>) and per blade (μmol O<sub>2</sub> h<sup>-1</sup> blade<sup>-1</sup>); NP<sub>max</sub> = maximum net photosynthesis per area (μmol O<sub>2</sub> cm<sup>-2</sup> h<sup>-1</sup>) and per blade (μmol O<sub>2</sub> h<sup>-1</sup> blade<sup>-1</sup>); R<sub>D</sub> = Dark respiration (μmol O<sub>2</sub> cm<sup>-2</sup> h<sup>-1</sup>); R<sub>L</sub> = light-enhanced respiration (μmol O<sub>2</sub> cm<sup>-2</sup> h<sup>-1</sup>); α = photosynthetic efficiency (μmol O<sub>2</sub> cm<sup>-2</sup> h<sup>-1</sup>/μmol incident quanta m<sup>-2</sup> s<sup>-1</sup>); E<sub>c</sub> and E<sub>k</sub> = compensation and saturation irradiances (μmol quanta m<sup>-2</sup> s<sup>-1</sup>); Φ<sub>max</sub> = Quantum yield (mol O<sub>2</sub> released mol photons absorbed<sup>-1</sup>); I/Φ<sub>max</sub> = minimum quantum requirements (photons required to release one O<sub>2</sub> molecule)

No differences in the light absorption efficiency of the photosynthetic pigments, quantified as a*_PAR_ and a*_Chl*a*_, were detected among variants (**Table 1**). However, when comparing values for all samples, some patterns emerged. *S. natans* I presented a significant linear and positive association between A_Chl*a*_ and Chl*a* (**Fig. 3B)** and a negative, nonlinear, association between a*_Chl*a*_ and Chl*a* (**Fig. 3C**; SI-Table S4). Negative, nonlinear associations were also found between a*_PAR_ and Chl[*a+c*] (**Fig. 3D**) with significant differences among variants for the intercepts (ANCOVA-tests; SI-Table S5). These results indicate that chlorophylls are more efficient at absorbing light within the blades of *S. natans* I than within the blades of the other two types. To better illustrate these differences, we compared a*_Chl*a*_ in thalli with intermediate Chl*a* contents (20-30 mg Chl*a* m^-2^), as well as a*_PAR_ in highly pigmented thalli (>38 mg Chl[a+c] m^-2^) of the two variants with that pigmentation. These comparisons uncovered significant differences in a*_Chl*a*_ between *S. fluitans* and *S. natans* (III) (F=4.59; *p*=0.02; **Fig. 3E**), and in a*_PAR_ between *S. fluitans* and *S. natans* I (t-test=2.92; *p*=0.02; **Fig. 3F**).

### Response to light

Photosynthesis–Irradiance curves (P-E curves) determined at 28°C unveiled significant differences among *Sargassum* variants when normalized either to area, weight, or blade size (**Fig. 4**; **Table 1**). The maximum net (NP_max_) and gross (GP_max_) photosynthetic rates of the two *S. natans* types were similar and significantly greater than the rates of *S. fluitans*. *S. natans* VIII enhanced photosynthesis production per blade by 4.5 times with respect to *S. natans* I and *S. fluitans* (**Fig. 4C, F**). Under low-subsaturating light conditions, *S. natans* I presented the largest productivity (highest photosynthetic efficiency, α; **Fig. 4I**; **Table 1**). Light-enhanced respiration, as expected, was greater than dark respiration (R_L_ > R_D_), and both rates significantly varied among algal types. The highest rates were determined for *S. natans* I, *S. fluitans* presented the smallest and *S. natans* VIII intermediate values (**Fig. 4J**). Notably, thallus respiration strongly contributed to the differences found in net photosynthesis. *S. natans* I, the species with the highest metabolic rates and smallest blade size, presented lower net photosynthetic production at 28°C than *S. natans* VIII, and the lowest carbon balances (P_max_:R_L_) (**Fig. 4K**). Indeed, *S. natans* VIII achieved the greatest production at that temperature thanks to its reduced respiration rates (**Fig. 4G**), and *S. fluitans* was able to achieve P_max_:R_L_ >3.5, similar to those of *S. natans* VIII and significantly above those of *S. natans* I (≈2; **Fig. 4K**; **Table 1**), thanks to its lowest blade respiration.

**Figure 4.**
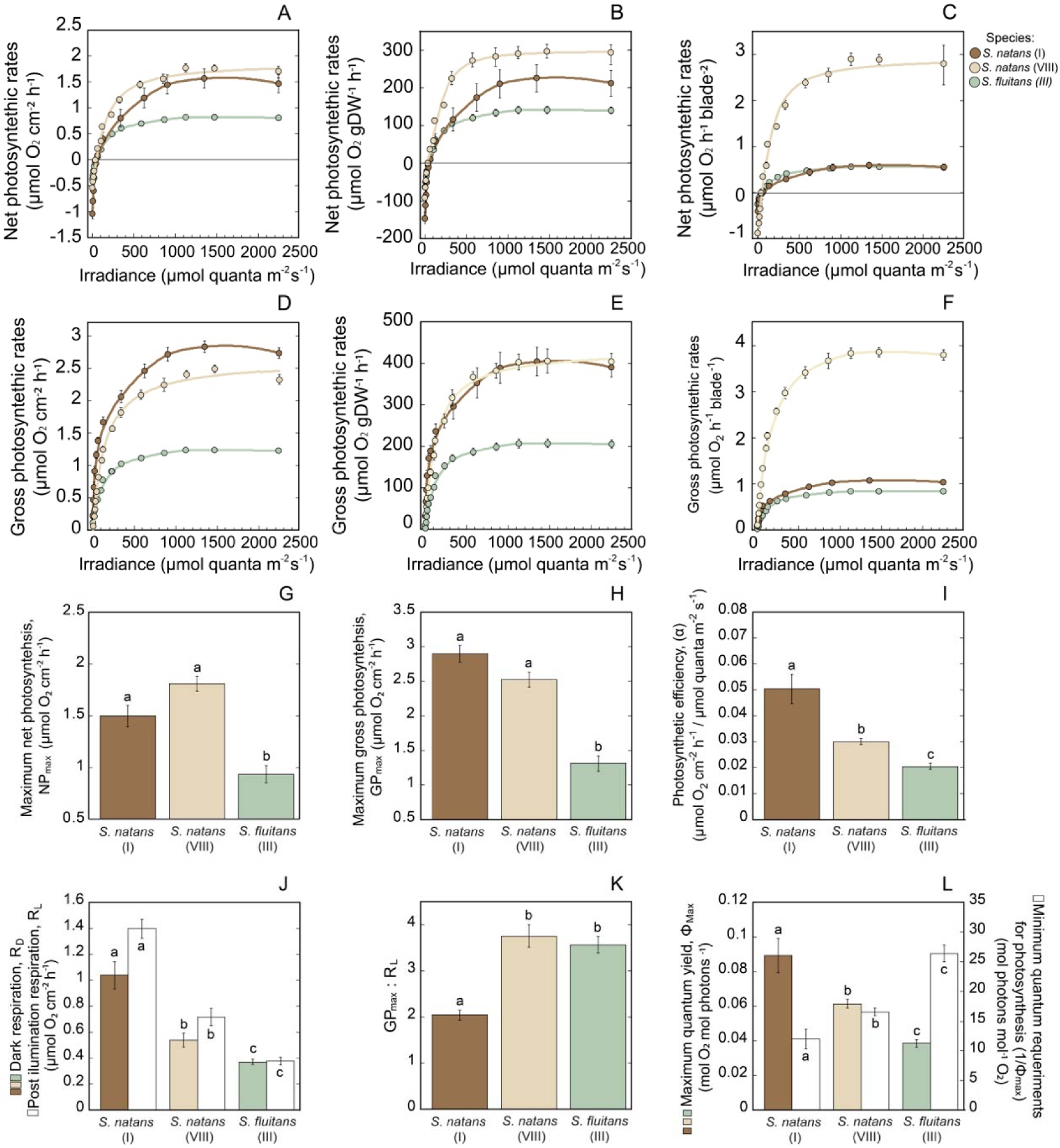
Light response of holopelagic *Sargassum*. Photosynthesis-Irradiance (P-E) curves for net (NP) and gross (GP) photosynthesis rates referred, respectively, to (A, D) area, (B, E) dry weight, and (C, F) thallus blade. In G-L plots are shown average ± SE values for (G) maximum net photosynthesis, NP_max_; (H) gross photosynthesis, GP_max_; and (I) the photosynthetic efficiency, α, for each genetic variant, together with the variation in (J) dark, R_D_, and light-enhanced, R_L_, respiration; (K) photosynthetic carbon balances above saturation, GP_max_: R_L_; and (L) the estimated values for the maximum quantum yield (Φ*_max_*, mol O_2_ evolved per mol photons absorbed) and the minimum quantum requirements of photosynthesis (*1/*Φ*_max_*, mol photons required to release one O_2_ molecule). Different letters indicate significant differences at the < 0.05 level, (Tukey Post-Hoc test). Colors are maintained consistent to identify values for each genetic variant: dark brown for *S. natans* I, light brown for *S. natans* VIII, and green for *S. fluitans* III. In graphs A-F dots describe the average values ± SE for each light treatment. The solid line illustrates the fit to a photosynthetic function (non-rectangular or bent-hyperbola model). A grey line in plots A-C illustrates the condition where photosynthesis and respiration are compensated (no variation in oxygen concentration in the determination).

The saturation (E_k_) and compensation (E_c_) irradiances were low for the three *Sargassum* variants, similar to the values reported for species from deeper waters (Stiger-Pouvreau et al. 2023). Significant differences were only found for E_k_ (**Table 1**). The highest quantum efficiency of the photosynthetic process (Φ*_max_*; mol O_2_ evolved per mol quanta absorbed) and lowest photosynthesis light requirements (*1/*Φ*_max_*; quanta required to evolve one O_2_ molecule; **Fig. 4L**) were observed for *S. natans* I. Intermediate values were estimated for *S. natans* VIII, whereas *S. fluitans* presented the highest light requirements (**Table 1**).

### Response to temperature

The annual temperature oscillation of the West Caribbean (24°-30°C) +4°C above the regional maximum in Puerto Morelos (Scheufen et al. 2017) and -2°C to the minimum, was chosen to characterize the temperature response. Significant differences for the optima of photosynthesis were found: *S. natans* VIII presented the NP_max_ optimum at 28°C, *S. natans* I at 26°C, and *S. fluitans* at 30°C (**Fig. 5A**). An analysis of the independent variation in photosynthesis and respiration rates uncovered a different GP_max_ optimum for *S. natans* I at 32°C (**Fig. 5B**). Algal respiration was maximized at 34°C in all *Sargassum* variants (**Fig. 5C, D**), indicative that no physiological temperature tipping point occurred within the range investigated. However, the metabolic scaling factor, Q_10_, revealed a differential positive effect of temperature on algal metabolic rates (**Table 1**). The highest effect on respiration was found for *S. natans* I (Q_10_ ≈3 for R_D_ and R_L_), whereas the lowest was observed in *S. fluitans* (Q_10_≈1.7), the species that presented the highest Q_10_ for photosynthesis (Q_10_ ≈2.5). Intermediate sensitivity to temperature was observed in *S. natans* VIII for both metabolic processes (**Fig. 5**; **Table 1**), together with the highest photosynthetic production per blade at all temperatures investigated (**Fig. 5E, F**). This differential effect of temperature on algal metabolic rates resulted in large variation in the thallus carbon balances (GP_max_:R_L_). *S. natans* VIII had an optimum at 28°C, with the largest values between 22°C and 28°C; *S. fluitans* extended its optimum between 28°C and 30°C; and the lowest GP_max_:R_L_ were found in *S. natans* I at all temperatures investigated. Differences were particularly large above 28°C (**Fig. 5G**). At 34°C, the three variants presented GP_max_:R_L_ ≤1, suggesting a common upper temperature threshold for pelagic *Sargassum* growth. Similarly, a common minimum was found for *1/*Φ*_max_* (maximum for Φ*_max_*) at 30°C, which may indicate a physiological optimum of photosynthesis performance for holopelagic *Sargassum*. This optimality, however, does not assume a peak-optimum in algal growth and productivity at 30°C, as the costs of: i) thallus respiration; ii) repair and maintenance of photosynthetic activity; and iii) the construction and life span of the new growing biomass; are not taken into account. Finally, the results also indicate that *S. fluitans* has minimum light requirements between 25 and 50 mol quanta, significantly above the values estimated for both *S. natans* variants (<20 mol quanta; **Fig. 5H**) for the temperature investigated.

**Figure 5.**
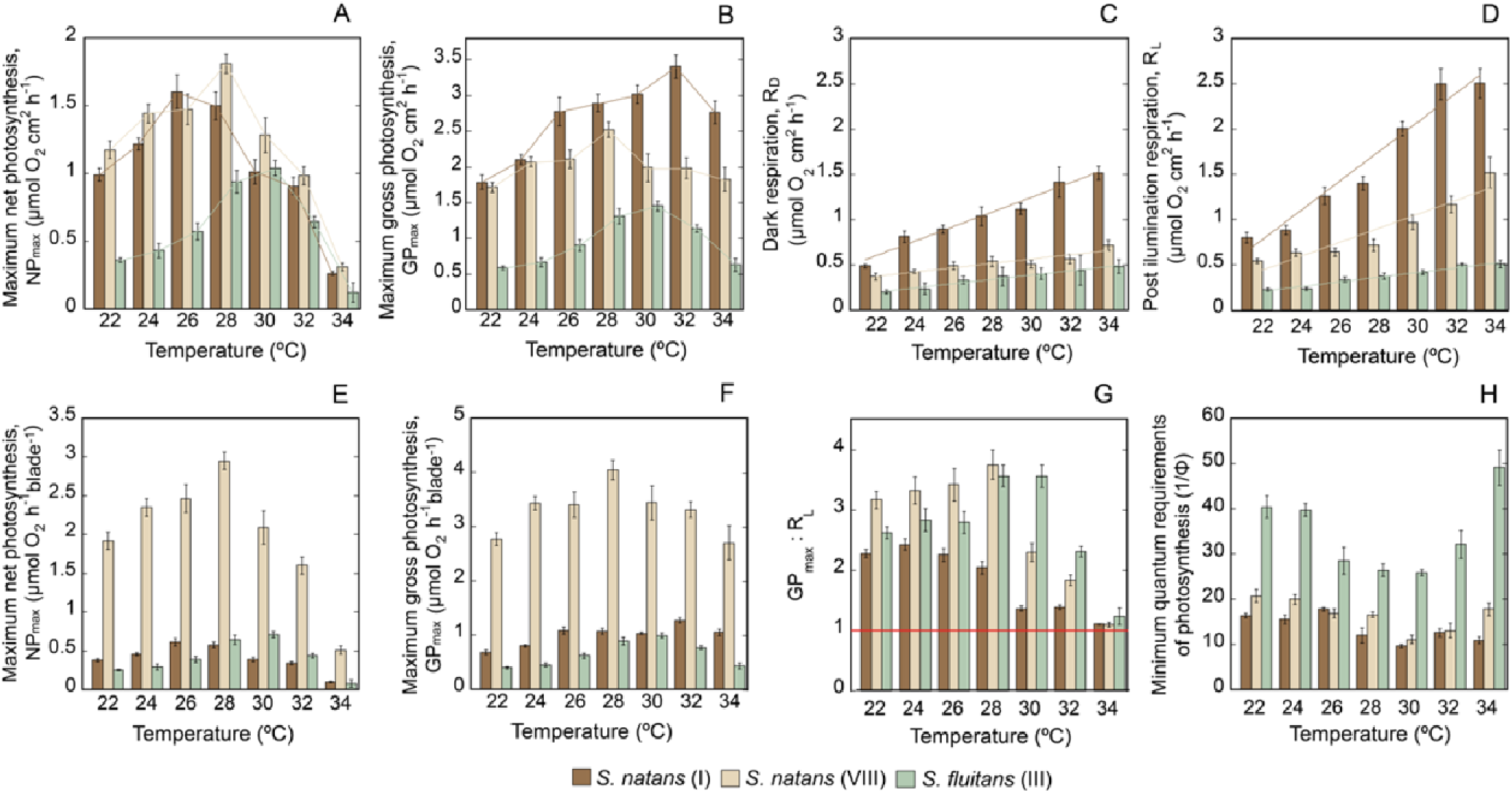
Temperature response of holopelagic *Sargassum*. Average ± SE for the variation of (A) maximum net photosynthesis, NP_max_; (B) maximum gross photosynthesis, GP_max_; (C) dark respiration, R_D_; (D) post-illumination (light enhanced) respiration, R_L_; (E-F) maximum net and gross photosynthesis per thallus bla e; (G) photosynthetic carbon balances (ratio of GP_max_: R_L_); and (H) minimum quantum requirements of photosynthesis at each temperature (1/Φ*_max_*, mol photons required to release one O_2_ molecule). Colors are maintained consistent for each variant: dark brown for *S. natans* I, light brown for *S. natans* VIII and green for *S. fluitans* III. Solid and fine lines illustrate the trend of variation of each parameter along the temperature range investigated. The red line in plot G indicates the carbon balance (GP_max_: R_L_ =1) that limits algal growth.

### Variation in biomass, size and growth rates

Literature data on the variation in the relative abundance of the *Sargassum* variants were obtained from 33 studies published between 1933 and 2022 (SI-Table S1). Data were examined for three large regions in the Atlantic: North Atlantic, West tropical Atlantic, and East tropical Atlantic (Africa); for the Caribbean and Gulf of Mexico; and for two periods, before and after 2011 **(Fig. 6A)**. Four regions in the Caribbean (West, Central, East and South) were also examined **(Fig. 6B)**. The literature concerning the variation in growth and blade size included, respectively, 10 studies published from 1983 to 2022 (SI-Table S2), and 9 studies with data collected from 2014 to 2020 (SI-Table S3). The first comparison revealed significant regional differences in variant dominance, and also before and after 2011 (**Fig. 6**). *S. fluitans* dominates in the West, Central and South of the Caribbean, the east tropical Atlantic (Africa), and the Gulf of Mexico, whereas *S. natans* VIII dominates in the small Antilles (East Caribbean) and West tropical Atlantic. Interestingly, *S. natans* I was predominant in the North Atlantic and Sargasso Sea before 2011 within the most diverse community ever characterized (**Fig. 6A**). However, after 2011, the two genetic variants that were better “adapted” to warmer temperatures also dominate in the North Atlantic algal community.

**Figure 6.**
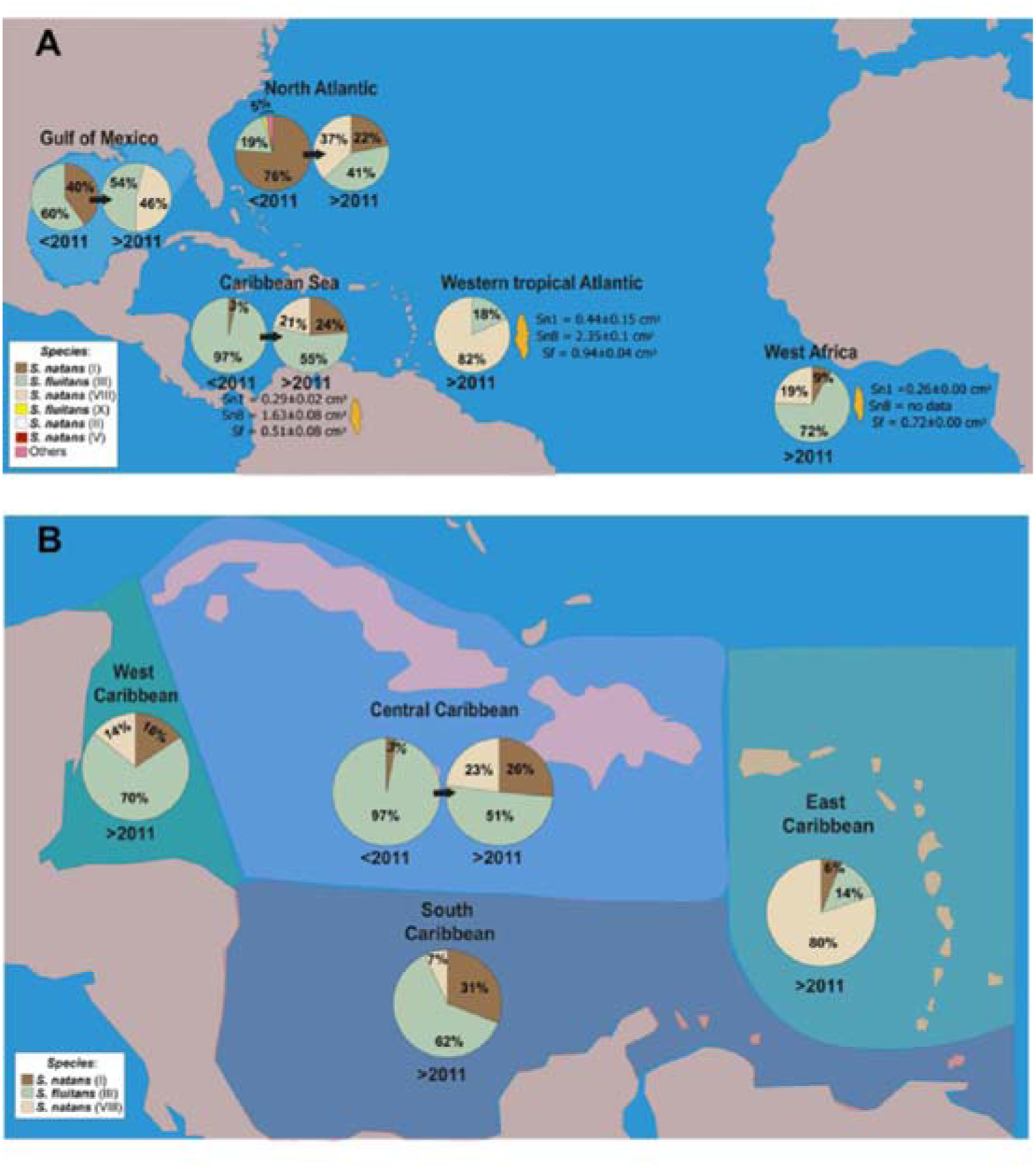
Spatial and temporal variability in relative abundance of the three genetic variants of holopelagic *Sargassum* in the Atlantic, Caribbean Sea and Gulf of Mexico. (A) Variation in relative abundance for the whole region before and after 2011, and changes, in orange, in blade size described as total projected blade area. (B) Variation in relative abundance for four regions of the Caribbean Sea, before and after 2011. Data and references from the literature are documented in: i) Table S1 of the supplementary information for changes in relative abundances of the variants; and ii) Table S3 for changes in blade size. Pies describe the variation in relative abundances among variants and before and after 2011.

Literature data for the relative growth rates (RGR, day^-1^; n=191) presented severe limitations for the comparability of data. Insufficient information on the field or experimental conditions under which RGR determinations were performed hampered this comparison. Moreover, we also identified large, suboptimal variability among replicates in a notable number of studies, and too many negative values for field or laboratory conditions that would be, according to our physiological characterizations, optimal for *Sargassum* growth. These problem constrained the analysis to the estimations of average RGR values for each variant, using only data determined under comparable conditions or treatments (**Fig. 7**; SI-Table S2). Significant differences were only detected between *S. natans* I and *S. fluitans,* as no differences were observed between the two *natans* variants (**Fig. 7**; SI-table S6). Average values suggest that the largest variant, *S. natans* VIII, presents the slowest growth rate, whereas *S. fluitans* is the species with the fastest growth, although this interpretation is not yet certain.

**Figure 7:**
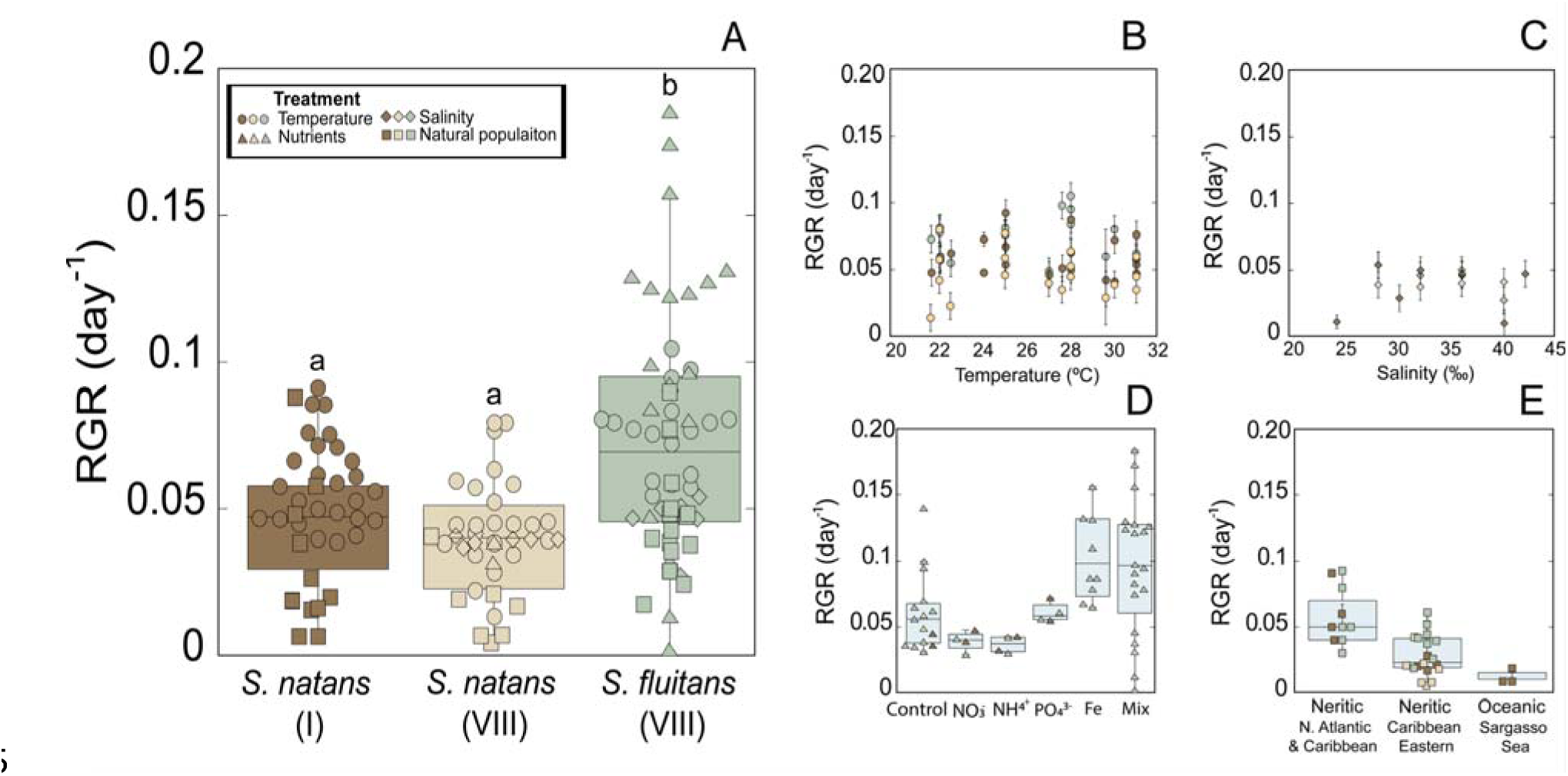
Variation in the relative growth rates of the three genetic variants of holopelagic *Sargassum*. (A) Box plots describing the variation of the Relative Growth Rates (RGR, day^-1^) reported in the literature for each genetic variant; and (B-E) RGR data reported for natural populations growing in oceanic conditions, and for experimental determinations on samples exposed to different treatments. Box plots encompass the 25% and 75% quartiles, and the central solid line indicates the median. Different symbols in plot A indicate the source of this information, the type of experimental conditions (B-D) or natural populations in neritic or oceanic conditions (E). Neritic refers to an oceanic region relatively shallow (<200 m depth), located just before the deep decline in the continental platform initiates. Data and references are documented in table S2 of the supplementary materials.

## Discussion

The optical and metabolic traits of pelagic *Sargassum* characterized corroborate and are in line with the genetic differences previously established between the three genetic variants of pelagic *Sargassum* present in the GASB (Amaral Zettler et al. 2016; González-Nieto et al. 2020; Siuda et al. 2024). We could document substantial differences in the light absorption capacity and efficiency of blades and photosynthetic pigments; in thallus photosynthesis and respiration rates; in the response to low light and minimum light requirements of photosynthesis; and in the effect of temperature on algal metabolic rates and photosynthetic production. *S. natans* I, the smallest variant, presented the highest light absorption capacity (A) and pigment absorption efficiency (a*); the highest metabolic rates (GP_max_, R_D_ and R_L_); photosynthetic (α) and quantum (Φ*_max_*) efficiencies; and the lowest light requirements (1/Φ*_max_*). This variant also showed the greatest phenotypic variation in thallus pigmentation and light absorption capacity, and was found as the most efficient and productive under resource limitations, not only light but also nutrient constraints, as its small size indicates lower carbon and nutrient requirements for biomass duplication. Moreover, its higher light absorption efficiency allows *S. natans* I to collect similar or even more solar energy than the other variants, with a more economical light harvesting system (i.e., lower pigmentation and nitrogen requirements). However, its net photosynthetic production and thallus carbon balance above saturation (GP_max_:R_L_) were below than those of *S. natans* VIII and *S. fluitans* at all temperatures investigated. The optimum carbon balance for *S. natans* I was determined at 24°C, consistent with the reported maximum growth rates (Hanisak and Samuel 1987; Corbin and Oxenford 2023), and despite the fact that the maximum photosynthesis, GP_max_, occurred at 32°C. These findings indicate that the lower photosynthetic performance of *S. natans* I at elevated temperature cannot be attributed to thermal stress but to its high respiratory demands, which reduce net carbon balances to values ≈1 or lower, negatively affecting photosynthetic production. In contrast, the observed reductions in thallus respiration in the other two variants allow enhancing their performance at elevated temperatures. Algal productivity in *S. natans* VIII increased between 22°C and 28°C, whereas *S. fluitans*, the species with the lowest metabolic rates, achieved the highest carbon balanced above 28°C. Therefore, this functional analysis can explain the dominance of *S. natans* VIII and *S. fluitans* in the warmer oceanic conditions of the NERR, which ranges under normal conditions from 28° to 31°C in summer (Marsh et al. 2023). *S. natans* VIII has a greater potential to dominate between 22°C and 28°C, and *S. fluitans* at 28°–30°C. A common threshold for pelagic *Sargassum* growth was also observed at 33°-34°C, which may explain why the great algal population documented for March 2023 (https://optics.marine.usf.edu/projects/saws.html) disappeared rapidly during one of the warmest summers so far recorded for the Northern Hemisphere (Esper et al. 2023). Our analysis can also explain the low abundance of *S. natans* I in the NERR, and why is a less competitive variant at lower latitudes, given its small size and high respiratory demands.

Literature data support part of these conclusions as *S. fluitans* dominates in the warmest regions (West, Central and South of the Caribbean and coastal regions of Africa) and also in summer (García-Sánchez et al. 2020), as predicted by its greater performance above 28°C, whereas *S. natans* I abundance in the GASB rises in the cold season (García-Sánchez et al. 2020). Remarkably, the largest interannual variability in the Mexican Caribbean was displayed by *S. natans* VIII (García-Sánchez et al. 2020; Vázquez-Delfín et al. 2021). *S. natans* I was also dominant in the North Atlantic before 2011, but it is not clear why this dominance was lost since 2011. However, we can explain why *S. natans* I still maintains its presence in the GASB, as the high light absorption, photosynthetic and quantum efficiencies of this variant, and its lowest light requirements support a greater ability of *S. natans* I to grow within dense canopies, a presence that will also improve when light, temperature and/or nutrient limitations may enhance its competitive ability with the other two variants.

The reductions found in thallus respiration and in the scaling metabolic factor, Q_10_, do reflect ‘adaptive’ traits of holopelagic *Sargassum* to warmer latitudes and seasons. Previous descriptions of mitochondrial *Sargassum* genomes (mtDNA; Amaral Zettler et al. 2016) have documented small genetic alterations, which involve relevant amino acid substitutions in encoded proteins for Complex I and Complex IV of the mitochondrial-oxidative-phosphorylation-system (OXPHOS). Although phylogenetic analyses generally assume the neutrality of mitochondrial genetic markers, this assumption has been questioned due to potential repercussions for energy production and organism’s life adaptation (Ballard and Pichaud 2014). Complex I is the first enzyme of the respiratory chain (NADH: ubiquinone oxidoreductase), fundamental for generating the proton-motive force across the inner mitochondrial membrane and a major contributor to the cellular production of reactive oxygen species (ROS), whereas Complex IV is the final and rate-limiting step of the respiratory chain (cytochrome c oxidase, COX), and then another key regulatory center of oxidative phosphorylation. Hence, although this analysis cannot support any mechanistic link between the physiological variability characterized and the genetic alterations documented (Amaral Zettler et al. 2016), it may stimulate future research to elucidate whether these mtDNA alterations are part of an adaptive response of pelagic *Sargassum* to heat, perhaps, through the contribution of metabolic tuning and epigenomics to fix asexual lineages. In this sense, this study underlines the potential of pelagic *Sargassum* for investigating how adaptive traits can be fixed in asexual lineages.

Overall, our analysis revealed that global warming favors the increasing presence of the two *Sargassum* variants best adapted to warmer temperatures in the Atlantic, but cannot support the hypotheses that global warming is the main driver of the *Sargassum* bloom, or why a once-rare variant such as *S. natans* VIII is currently the primary contributor to GASB in some areas. Indeed, as mentioned above, our results cannot explain why *S. natans* I dominated in the North Atlantic before 2011 in a more diverse community, and why *S. natans* VIII has become so abundant in the Atlantic since 2011. Nutrient enrichment could contribute to explain these uncertainties, as some *Sargassum* species can become invasive when nutrient availability increases (Stiger-Pouvreau et al. 2023). However, the maximum photosynthetic rates determined here for pelagic *Sargassum* are within the average values reported for macroalgae (≈2–6.4 mg O_2_ gDW^-1^; Enríquez et al. 1995). Moreover, literature data for relative growth rates showed average values significantly below the mean reported for a diverse group of macroalgae (Nielsen et al. 1996), and certainly well below the values determined for invasive *Sargassum* species (Stiger-Pouvreau et al. 2023). This indicates that the massive presence of pelagic *Sargassum* in the tropical Atlantic depends, primarily, on substantial improvements in nutrient availability at the ocean surface, capable of triggering and maintain a highly productive *Sargassum* bloom (Glibert et al. 2018). The vast amount of organic nutrients that have been estimated for *Sargassum* (Wang et al. 2018) is the strongest evidence for the fundamental role of nutrients in explaining GASB formation, since large amounts of inorganic nutrients must have been present in the ocean surface before they were assimilated by the algae and retained in their tissues. It has been recently proposed that equatorial upwelling of phosphorus drives *Sargassum* blooms (Jung et al. 2025) linked to the well-known N_2_ fixation capacity of the epiphytic diazotrophic community of pelagic *Sargassum* (e.g., Phlips and Zeman 1990; Johnson et al. 2023), a new hypothesis that does not rule out other possible sources of fertility in the tropical Atlantic such as nutrient fluxes from the Amazon and Orinoco Rivers in the western Atlantic, and the Congo River in the eastern equatorial recirculation area (Oviatt et al. 2019), and the potential contribution of iron fertilization from Saharan dust (Gouvêa et al. 2020; Hernández-Ayala and Méndez-Tejeda 2024). In fact, although the activation of algal blooms (Glibert et al. 2018), and the positive effect of nutrients on the growth (Lapointe 1986, 1995) and productivity (Lapointe 1986, 1995; Lapointe et al. 2014) of holopelagic *Sargassum* has to play a central role in GASB formation, our analysis supports the key contribution of global warming by improving the performance of two genetic variants in a warmer ocean. Therefore, nutrients cannot be considered the only driver of the massive presence of pelagic *Sargassum* in the NERR, highlighting the crucial contribution of a deeper understanding of pelagic *Sargassum* biology to fully explaining a biological phenomenon ultimately governed by biological rules.

Hence, the capacity of a warming ocean to supply the nutrients and energy (light) that invigorates *Sargassum* growth and its life span across the NERR is a plausible explanation for GASB formation. Moreover, nutrient fertilization in a warmer ocean could also explain the ecological success of *S. natans* VIII, a previously rare variant in the Atlantic, together with the loss of dominance of *S. natans* I in the northern region, an efficient variant under resource limitations but less competitive with greater nutrient availability and water temperatures above 22-24°C. Our study also revealed that *S. natans* VIII and *S. fluitans* are less efficient in the use of the available resources, suggesting a more opportunistic strategy for these variants. The largest size and, then, higher structural costs of *S. natans* VIII supports higher nutrient requirements and then lower performance for this variant under nutrient-limiting conditions, which reinforces the interpretation that the current ecological success of *S. natans* VIII in the Atlantic may respond to greater nitrogen availability in the ocean. On the other hand, the higher performance of *S. fluitans* above 28°C and a greater resistance to starvation could also explain its dominance in warmer regions, and its greater presence before 2011.

The critical question that remains to be resolved is where such huge amount of organic nutrients of the pelagic *Sargassum* community in the Atlantic come from. There is evidence that thallus nutrient content of pelagic *Sargassum* has increased significantly (Lapointe et al. 2021), with the highest contents documented for the western tropical Atlantic and eastern Caribbean (McGillicuddy et al. 2023). Phytoplankton blooms were also reported in the northeastern Caribbean two years before the massive population of floating *Sargassum* was detected in the tropical Atlantic (2009, 2010), events that had not been observed in the previous 30 years, and that were attributed to nutrient fluxes from the Amazon River plume (Johns et al. 2014). The literature comparison and our own determinations also revealed that the blades of the *Sargassum* populations growing in the Eastern Caribbean and Western Tropical Atlantic are larger (Schell et al. 2015; SI*-*Table S3) relative to specimens collected near the coasts of Nigeria (Oyesiku et al. 2014) or Mexico (Vásquez Elizondo et al. 2023; and this study). Both observations, phytoplankton blooms and blade size (Tanaka et al. 2022) suggest higher nutrient availability for *Sargassum* growth in the Western tropical Atlantic (Tanaka et al. 2022). Furthermore, the abnormally high arsenic concentration reported for *Sargassum* tissues (Ortega-Flores et al. 2022; McGillicuddy et al. 2023; Bousso et al. 2024), with the highest contents also found for the populations of the Western tropical Atlantic (Bousso et al. 2024), provides a confident tracer for the main nutrient sources that feed the *Sargassum* blooms. The concentration of arsenic in the surface of the Atlantic is relatively low (≈ 1.7 μg/L, Neff 1997) and, therefore, insufficient to explain that high reported content, and arsenic uptake by *Sargassum* is facilitated when phosphorus is absorbed from the water, as it is mediated by phosphate transporters, which confound it with phosphate (Devault et al. 2020). Accordingly, a reasonable hypothesis to explain the high arsenic concentration in *Sargassum* tissues is that a significant fraction of algal growth occurs in an area rich in both elements, arsenic and phosphorus. High concentration of arsenic has been reported for the Amazonian Fan (Sullivan and Aller 1996; Scarpelli 2005) due to mining activities in the Andes. Hence, the fact that the *Sargassum* populations of the Western tropical Atlantic (Bousso et al. 2024) present the greatest arsenic concentration in their tissues, highest nutrient contents, and larger blades strengthens the interpretation that the primary sources of nutrients that feed the GASB (mainly phosphorus) are located where two large rivers originating in the Andes, Orinoco and Amazon, release their waters enriched in both arsenic and phosphorus. Arsenic bioaccumulates in *Sargassum* tissues in the form of various organo-arsenic compounds, explaining its increasing concentration with respect to phosphorus (McGillicuddy et al. 2023), and element that is rapidly metabolized and depleted from *Sargassum* growing tissues, especially in areas with lower phosphorus availability.

Two areas of *Sargassum* aggregations have been observed at the initiation of the growing season (i.e., March, April, and May), exactly where the Orinoco and Amazon Rivers plumes are located (Jouanno et al. 2023). Nutrients are removed by the algae during the growing season, which sets limits to annual algal production until new nutrient fluxes can reactivate algal blooms. So, the interannual variability of *Sargassum* biomass may be controlled by the fluctuation of nutrient fluxes from the coast, and algal growth and nutrient removal in the preceding season, although the contribution of phosphorus upwelling to this seasonal, interannual and spatial variability also needs to be investigated (Jung et al. 2025). The two *Sargassum* aggregation documented by Jouanno et al. (2023) in March-April-May may reveal the location of two important *Sargassum* blooms, and/or where the new growing algae are primarily aggregated, as oceanic currents may mask the location of the blooms by aggregating or spreading the new biomass towards different oceanic routes. *Sargassum* populations will maintain a variable growth and biomass dynamics in its community during transport in regions with different light and temperature regimes, as well as reduced nutrient availability outside of nutrient-enriched areas. Our study has characterized the photosynthetic physiology of the *Sargassum* populations that reach the northern Mexican Caribbean. While the relative differences between variants may not vary, absolute values are expected to be higher near the bloom, similarly to the enhancement observed in blade size for the populations of the western tropical Atlantic (Schell et al. 2015). Other biological and oceanic/climatic processes need to be elucidated to fully undersand GASB variability such as: i) determinations of more reliable values for algal growth and thallus lifespan under different light, temperature, and nutrient regimes; ii) understanding the contribution of oceanic currents and climatic events to the aggregation or dispersal of *Sargassum* populations in different ocean regions; and iii) the role of *Sargassum* canopy in facilitating algal growth and/or extending thallus life span.

A putative role of the canopy was unveiled by the low light requirements, and compensation (E_C_) and saturation (E_K_) irradiances determined in this study for the three *Sargassum* variants, surprisingly low for species ‘adapted’ to live on the ocean surface (Stiger-Pouvreau et al. 2023). This suggests high costs of living of pelagic *Sargassum* at full solar irradiance (Aro et al. 1993; Raven 2011; López-Londoño et al. 2022), highlighting a potential role of *Sargassum* canopy in abating these costs, similarly to the photoprotective role documented for seagrass canopies (Schubert et al. 2015). The combination of high photosynthetic and quantum efficiencies with the ability to form dense canopies may represent a key achievement of pelagic *Sargassum* to allow dense patches to enhance photosynthetic production, highlighting a potential positive feedback in algal production that needs to be understood. Dense *Sargassum* canopies can be formed by several variants or by monospecific populations, as those reported for *S. fluitans* near Cuba (Azanza-Ricardo and Pérez-Martín 2016) and Nigeria (Adet et al. 2017).

The massive presence of holopelagic *Sargassum* in the tropical Atlantic illustrates the Anthropocene capacity to transform not only coastal ecosystems but the entire ocean in ways that favor the growth of opportunistic/invasive algae. Predictive models for the variation of S*argassum* biomass will enhance the understanding of the mechanisms that govern *Sargassum* growth and the persistence of massive algal populations in the Atlantic. The optical and physiological traits documented here can aid in their development, paving the way for the development of effective solutions to contend this serious environmental crisis on the coast. These models may help to explain, for instance, why in the year with the world’s warmest temperatures on record in summer (https://www.noaa.gov/news/2023-was-worlds-warmest-year-on-record-by-far), a large fraction of the largest ever recorded *Sargassum* biomass in the tropical Atlantic, March 2023, disappeared during the warmest months (https://optics.marine.usf.edu/projects/saws.html), and, therefore, the alarming golden tides predicted for the Caribbean that year never arrived. Our analysis found at 34°C a temperature threshold, indicative that when the tolerance limit of *Sargassum* physiology is surpassed, global warming can also limit the expansion of this alga in the Atlantic, particularly during the hottest season. The control of such massive algal presence in the Atlantic is urgent as the massive *Sargassum* tides that result when these populations arrive to the coast affect not only beaches with touristic economic interest but also entire coastal ecosystems in the Caribbean, northern South America, West Africa, and the Gulf of Mexico, where coral reefs are particularly threatened by coastal eutrophication (Velázquez-Ochoa and Enríquez 2023). The determination of ‘when’ and ‘where’ *Sargassum* has to be harvested in the ocean, as a major operation, is the only effective solution for controlling the blooms at the beginning of the growing season, and the environmental and economic impacts that the massive algal tides are causing since 2011. No realistic solutions can be implemented once *Sargassum* arrives massively on the coast.

## Acknowledgments

Authors acknowledge the contribution of Román M. Vásquez-Elizondo to perform some of the optical and physiological characterizations of *Sargassum fluitans*.

## Funding

The remaining funds of the European collaborative project (PFP7-FORCE-244161) to SE are acknowledged, as they support one year of technical assistance of Román M. Vásquez-Elizondo. Personal funds of SE support the development of this study. We also acknowledge the financial support of SECIHTI, former CONAHCYT, to RV-O, within the SNII-III economical support awarded to SE

## Competing interests

Authors declare that they have no competing interests.

## Author contributions

Conceptualization, Supervision, funding acquisition and Writing – review & editing: SE; Investigation: RV-O; Formal analysis, and Writing original draft: RV-O and SE

## Data availability

All data are available in the main text or in the supporting materials.

## Supporting Information (brief legends)

The PDF file includes: Tables S1 to S6 and 44 references for the literature data compilation.

**Table S1**: Literature data for the variation in community composition (i.e., relative abundance) of holopelagic Sargassum throughout the Atlantic Ocean and the Caribbean.

**Table S2**: Literature data for relative growth rates (RGR) estimations for the three genetic variants of holopelagic Sargassum present in the GASB.

**Table S3**: Literature data for the variation in thallus size (blade area) of the three holopelagic Sargassum variants present in the GASB.

**Table S4**: Linear and non-linear (power-functions) for the associations of variation between optical and structural traits of holopelagic Sargassum.

**Table S5**: ANCOVA tests for investigating significant differences in the associations of variation between optical traits and pigment content per projected area within each variant.

**Table S6**: Analysis of variance (ANOVA tests) for investigating differences in the relative growth rates (RGR) values documented in the literature, for the three holopelagic Sargassum variants present in the Central tropical Atlantic.

## Notes

### Competing Interest Statement

The authors have declared no competing interest.

