## Supplemental Tables S1-S6 for "Adaptation to heat and ocean fertilization, two keys for understanding the massive *Sargassum* growth in the Atlantic"

**The PDF file includes:**

Tables S1 to S6 and 44 references for the literature data compilation.

**Table S1:** Literature data for the variation in community composition (i.e., relative abundance) of holopelagic *Sargassum* throughout the Atlantic Ocean and the Caribbean.

**Table S2**: Literature data for relative growth rates (RGR) estimations for the three genetic variants of holopelagic *Sargassum* present in the GASB.

**Table S3**: Literature data for the variation in thallus size (blade area) of the three holopelagic *Sargassum* variants present in the GASB.

**Table S4:** Linear and non-linear (power-functions) for the associations of variation between optical and structural traits of holopelagic *Sargassum*.

**Table S5:** ANCOVA tests for investigating significant differences in the associations of variation between optical traits and pigment content per projected area within each variant.

**Table S6:** Analysis of variance (ANOVA tests) for investigating differences in the relative growth rates (RGR) values documented in the literature, for the three holopelagic *Sargassum* variants present in the Central tropical Atlantic.

**Table S1.** Literature data for the variation in community composition (i.e., relative abundance) of holopelagic *Sargassum* throughout the Atlantic Ocean and the Caribbean.

| **Sampling date** | **Region** | **Locality** | **Species/variant composition (% cover)** | | **Literature Source** |
|---|---|---|---|---|---|
| 1933-1935 | North Atlantic | Sargasso Sea | *S. natans* (I) | 72.1% | 1 |
| | | | *S. fluitans* (III) | 17.3% | |
| | | | *S. natans* (VIII) | 0% | |
| | | | *S. fluitans* (X) | 7.7% | |
| | | | *S. natans* (II) | 2.88% | |
| 2-15 October, 1962 | North Atlantic | Sargasso Sea gyre (From east of Colon, C. Z. to the offing of Block Island Sound, Rhode Island) | *S. natans* (I) | 80% | 2 |
| | | | *S. fluitans* (III) | 10% | |
| | | | The remaining 10% was not explained. | | |
| 1966, 1969, 1971-1975 | North Atlantic | Sargasso Sea, Gulf Stream, and south of the subtropical convergence zone | *S. natans* (I) | > 95% | 3 |
| | | | Rest of species | < 5% | |
| Mar, 1979 | North Atlantic | Sargasso Sea: northern and southern regions (207 oceanic stations) | *S. natans* (I) | 80 – 90% | 4 |
| | | | *S. fluitans* (III) | 15 – 23% | |
| | | | *S. natans* (VIII) | 0% | |
| | | | *S. natans* (V) | 2 – 5% | |
| | | | *S. natans* (II) | 2 – 5% | |
| July, 1981 | North Atlantic | Sargasso Sea and Gulf Stream | *S. natans* (I) | 54% | 5 |
| | | | *S. fluitans* | 46% | |
| | | | *S. natans* (VIII) | 0% | |
| From March to December of 1982 | North Atlantic | Bermuda, an oceanic island in the Sargasso Sea nearly 1,000 km from nearest land | *S. natans (I)* | > 60% | 6 |
| | | | *S. fluitans* | < 40% | |
| | | | *S. natans* (VIII) | 0% | |
| 2011-2012 | North Atlantic | Sargasso Sea: from Bermuda in the north to the Bahama Islands in the south | *S. natans* (I) | > 95% | 3 |
| | | | Rest of species | < 5% | |
| 2015-2016 | North Atlantic | Sargasso Sea | *S. natans* (I) | 43.5% | 7 |
| | | | *S. fluitans* (III) | 34% | |
| | | | *S. natans* (VIII) | 22.5% | |
| November 2015 to May 2015 | North Atlantic | South Sargasso Sea | *S. natans (I)* | *87.5%* | 8 |
| | | | *S. fluitans* (III) | ~6% | |
| | | | *S. natans (VIII)* | ~6% | |
| 2015-2018 | North Atlantic | North Sargasso Sea | *S. natans (I)* | 0% | 9 |
| | | | *S. fluitans* (III) | ~80% | |
| | | | *S. natans* (VIII) | ~20% | |
| 2015-2018 | | South Sargasso Sea | *S. natans* (I) | 0% | 9 |

| | | | | | |
|---|---|---|---|---|---|
| | North Atlantic | | *S. fluitans* (III)<br>*S. natans* (VIII) | ~70%<br>~30% | |
| 2015-2016 | North Atlantic | Gulf Stream | *S. natans* (I)<br>*S. fluitans* (III)<br>*S. natans* (VIII) | 13.6%<br>0%<br>86.3% | 7 |
| 2015-2018 | North Atlantic | North Gulf Stream | *S. natans* (I)<br>*S. fluitans* (III)<br>*S. natans* (VIII) | 0%<br>0%<br>100% | 9 |
| 2015-2018 | North Atlantic | South Gulf Stream | *S. natans* (I)<br>*S. fluitans* (III)<br>*S. natans* (VIII) | 0%<br>100%<br>0% | 9 |
| May 2000 | Gulf of Mexico | Northwestern Gulf of Mexico: region of the United States | *S. natans* (I)<br>*S. fluitans* (III) | 40%<br>60% | 10 |
| 2015-2016 | Gulf of Mexico | Gulf of Mexico | *S. natans* (I)<br>*S. fluitans* (III)<br>*S. natans* (VIII) | 0%<br>53.6%<br>46.4% | 7 |
| 1993-1997 | Caribbean Sea<br>(Central Caribbean) | South coast of Cuba | *S. natans (I)*<br>*S. fluitans* (III)<br>*S. natans* (VIII) | 3%<br>97%<br>0% | 11 |
| May 2012 | Caribbean Sea<br>(Central Caribbean) | Central-south coast of Cuba | *S. natans* (I)<br>*S. fluitans* (III)<br>*S. natans* (VIII) | 0%<br>100%<br>0% | 12 |
| June-July 2015 | Caribbean Sea<br>(Central Caribbean) | Guanahacabibes peninsula, Cuba | *S. natans* (I)<br>*S. fluitans* (III)<br>*S. natans* (VIII) | 0%<br>100%<br>0% | 13 |
| May 2018 to May 2019 | Caribbean Sea<br>(Central Caribbean) | Eastern coast of Cuba | *S. natans* (I)<br>*S. fluitans* (III)<br>*S. natans* (VIII) | 45%<br>30%<br>25% | 14 |
| March 2019 | Caribbean Sea<br>(Central Caribbean) | North coast of Cuba | *S. natans*<br>*S. fluitans* (III) | < 40%<br>> 60% | 15 |
| June 2019 to June 2021 | Caribbean Sea<br>(Central Caribbean) | La Habana, Cuba | *S. natans* (I) ><br>*S. fluitans* (III), *S. natans* (VIII) | | 16 |
| September 2019 to March 2020 | Caribbean Sea<br>(Central Caribbean) | Northwest coast of Cuba | *S. natans* (I)<br>*S. fluitans* (III)<br>*S. natans* (VIII) | 41 – 63%<br>25 – 36%<br>12 – 31% | 17 |
| May 2020 to August 2021 | Caribbean Sea<br>(Central Caribbean) | Northwest coast of Cuba | *S. natans* (I)<br>*S. fluitans* (III)<br>*S. natans* (VIII) | 5 – 12%<br>52 – 56%<br>33 – 43% | 17 |
| March to April 2021 | Caribbean Sea<br>(Central Caribbean) | Northwest coast of Cuba | *S. natans* (I)<br>*S. fluitans* (III)<br>*S. natans* (VIII) | 41 – 63%<br>25 – 36%<br>12 – 31% | 17 |

| | | | | | |
|---|---|---|---|---|---|
| 2018 | Caribbean Sea<br>(Central Caribbean) | Between Cuba and Dominican Republic | *S. natans* (I)<br>*S. fluitans* (III)<br>*S. natans* (VIII) | 0%<br>0%<br>100% | 9 |
| October 2015 | Caribbean Sea<br>(Central Caribbean) | Santo Domingo and Guayacanes, Dominican Republic | *S. natans*<br>*S. fluitans*<br>*S. natans* (VIII) | < 40%<br>> 60%<br>0 | 18 |
| 2015-2016 | Caribbean Sea<br>(Central Caribbean) | Central Caribbean | *S. natans* (I)<br>*S. fluitans* (III)<br>*S. natans* (VIII) | 2.6%<br>29%<br>68.4% | 7 |
| July 2020 | Caribbean Sea<br>(Central Caribbean) | Fort Rocky beach, Port Royal, Manchioneal and Port Maria, Jamaica | *S. natans* (I)<br>*S. fluitans* (III)<br>*S. natans* (VIII) | 8 – 28%<br>67 – 90%<br>2 – 5% | 19 |
| November 2015 to May 2015 | Caribbean Sea<br>(Eastern Caribbean) | Antilles current | *S. natans (I)*<br>*S. fluitans* (III)<br>*S. natans (VIII)* | 0%<br>8%<br>92% | 8 |
| November 2015 to May 2015 | Caribbean Sea<br>(Eastern Caribbean) | Islands from the Eastern Caribbean | *S. natans (I)*<br>*S. fluitans* (III)<br>*S. natans (VIII)* | 3.7%<br>1%<br>95.3% | 8 |
| December 2021 –February 2022 | Caribbean Sea<br>(Eastern Caribbean) | Morgan Lewis beach, Barbados | *S. natans* (I)<br>*S. fluitans* (III)<br>*S. natans* (VIII) | 8 – 11%<br>25 – 61%<br>29 – 67% | 20 |
| March to May 2022 | Caribbean Sea<br>(Eastern Caribbean) | Morgan Lewis beach, Barbados | *S. natans* (I)<br>*S. fluitans* (III)<br>*S. natans* (VIII) | 14 – 33%<br>43 – 55%<br>12 – 41% | 20 |
| June to August 2022 | Caribbean Sea<br>(Eastern Caribbean) | Morgan Lewis beach, Barbados | *S. natans* (I)<br>*S. fluitans* (III)<br>*S. natans* (VIII) | 19.5 – 26%<br>58 – 67%<br>12.5 – 18% | 20 |
| September to November 2022 | Caribbean Sea<br>(Eastern Caribbean) | Morgan Lewis beach, Barbados | *S. natans* (I)<br>*S. fluitans* (III)<br>*S. natans* (VIII) | 0 – 32%<br>8.5 – 60%<br>23 – 91.5% | 20 |
| September to November 2016 | Caribbean Sea<br>(Western Caribbean) | Puerto Morelos, México | *S. natans* (I)<br>*S. fluitans* (III)<br>*S. natans* (VIII) | 0%<br>35 – 100%<br>0 –60% | 21 |
| December 2016 to February 2017 | Caribbean Sea<br>(Western Caribbean) | Puerto Morelos, México | *S. natans* (I)<br>*S. fluitans* (III)<br>*S. natans* (VIII) | 0%<br>44 – 92%<br>8 – 56% | 21 |
| March to May 2017 | Caribbean Sea<br>(Western Caribbean) | Puerto Morelos, México | *S. natans* (I)<br>*S. fluitans* (III)<br>*S. natans* (VIII) | 0%<br>76 – 94%<br>6 – 24% | 21 |

| | | | | | |
|---|---|---|---|---|---|
| February to May 2018 | Caribbean Sea (Western Caribbean) | Puerto Morelos, México | *S. natans* (I)<br>*S. fluitans* (III)<br>*S. natans* (VIII) | 8 – 38%<br>57 – 83%<br>3 – 24% | 21 |
| June to August 2018 | Caribbean Sea (Western Caribbean) | Puerto Morelos, México | *S. natans* (I)<br>*S. fluitans* (III)<br>*S. natans* (VIII) | 9 – 40%<br>57 – 88%<br>1 – 5% | 21 |
| September to November 2018 | Caribbean Sea (Western Caribbean) | Puerto Morelos, México | *S. natans* (I)<br>*S. fluitans* (III)<br>*S. natans* (VIII) | 3 – 42%<br>50 – 95%<br>2 – 9% | 21 |
| September 2018 | Caribbean Sea (Western Caribbean) | From region northern to center: Cancún, Puerto Morelos, Xcalacoco, México | *S. natans* (I)<br>*S. fluitans* (III)<br>*S. natans* (VIII) | 6 – 35.9%<br>41 – 91%<br>1.2 – 3.6% | 22 |
| September 2018 | Caribbean Sea (Western Caribbean) | From center to southern region: Tulum and Playa Blanca, México | *S. natans* (I)<br>*S. fluitans* (III)<br>*S. natans* (VIII) | 6 – 17.9%<br>81.6 –91%<br>0.1 – 0.07% | 22 |
| December 2018 to February 2019 | Caribbean Sea (Western Caribbean) | Puerto Morelos, México | *S. natans* (I)<br>*S. fluitans* (III)<br>*S. natans* (VIII) | 28 – 46%<br>39 – 70%<br>3 – 10% | 21 |
| March to May 2019 | Caribbean Sea (Western Caribbean) | Puerto Morelos, México | *S. natans* (I)<br>*S. fluitans* (III)<br>*S. natans* (VIII) | 26 – 48%<br>41 – 73%<br>1 – 11% | 21 |
| June to August 2019 | Caribbean Sea (Western Caribbean) | Puerto Morelos, México | *S. natans* (I)<br>*S. fluitans* (III)<br>*S. natans* (VIII) | 13 – 28%<br>61 – 87%<br>0 – 3% | 21 |
| February to March 2019 | Caribbean Sea (Western Caribbean) | Puerto Morelos, México | *S. natans* (I)<br>*S. fluitans* (III)<br>*S. natans* (VIII) | 0 – 2%<br>36 – 88%<br>10 – 64% | 21 |
| April to May 2019 | Caribbean Sea (Western Caribbean) | Puerto Morelos, México | *S. natans* (I)<br>*S. fluitans* (III)<br>*S. natans* (VIII) | 0 – 14%<br>0 – 98%<br>2 – 100% | 21 |
| February to April 2019 | Caribbean Sea (Western Caribbean) | Puerto Morelos, México | *S. natans* (I)<br>*S. fluitans* (III)<br>*S. natans* (VIII) | 26%<br>65%<br>9% | 23 |
| September 2014 | Caribbean Sea (South Caribbean) | Northeastern coast of San Andres Island, Colombia | *S. natans* (I)<br>*S. fluitans* (III)<br>*S. natans* (VIII) | < 40%<br>> 60%<br>0% | 24 |
| March to April 2019 | Caribbean Sea (South Caribbean) | Cahuita, Puerto Vargas, Puerto Viejo, Playa Chiquita, Punta Uva, Manzanillo, Costa Rica | *S. natans* (I)<br>*S. fluitans* (III)<br>*S. natans* (VIII) | 20%<br>10%<br>65% | 25 |

| | | | | | |
|---|---|---|---|---|---|
| November 2015 to May 2015 | Tropical North Atlantic | North-west tropical Atlantic | *S. natans (I)*<br>*S. fluitans* (III)<br>*S. natans* (VIII) | 0%<br>12.7%<br>87.3% | 8 |
| 2015 | Tropical North Atlantic | Western tropical Atlantic | *S. natans* (I)<br>*S. fluitans* (III)<br>*S. natans* (VIII) | 0%<br>0%<br>100% | 9 |
| 2015-2016 | Tropical North Atlantic | Tropical Atlantic | *S. natans* (I)<br>*S. fluitans* (III)<br>*S. natans* (VIII) | 0%<br>41.4%<br>58.6% | 7 |
| 2011 | West Africa | Sierra Leone | *S. natans* (I)<br>*S. fluitans* (III)<br>*S. natans* (VIII) | 0%<br>100%<br>0% | 26 |
| May 2011 to august 2012 | West Africa | West Africa coast | *S. natans* (I)<br>*S. fluitans* (III)<br>*S. natans* (VIII) | 0%<br>100%<br>0% | 27 |
| May – June 2017 | West Africa | Western Nigerian coast | *S. natans* (I)<br>*S. fluitans* (III)<br>*S. natans* (VIII) | 0%<br>100%<br>0% | 28 |
| June 2016 | West Africa | Côte d’Ivoire | *S. natans* (I)<br>*S. fluitans* (III)<br>*S. natans* (VIII) | 0%<br>21.67%<br>78.33% | 29 |
| July, 2011 | Brazilian | Northern Brazilian coast | No difference quantified between<br>*S.fluitans and S. natans* | | 30 |

**Table S2.** Literature data for relative growth rates (RGR) estimations for the three genetic variants of holopelagic *Sargassum* present in the GASB.

| Objective | Treatment | Environmental conditions: TºC, light and salinity levels | Time incubation | Initial weight | No. replicates | Location | Specie | RGR ($d^{-1}$) | Source Ref. |
|---|---|---|---|---|---|---|---|---|---|
| Response to light | 13 light levels ranging from 8–576 | **Two experiment replicates. Light levels:** from 8 to 576 µE $m^{-2}$ $s^{-1}$**;** **temperature:** no specific; **salinity** 36‰ | 21 days | 20 mm of length | 6 replicates (3 in each experiment) | Neritic region (Florida Waters) | *S. fluitans* | 0.109 ± 0.002 | 31 |
| | Three light treatments: using a 12:12 h light: dark photoperiod. | **Two experiment replicates:** August-September of (1) 2019 and (2) 2020. **Light levels:** Low:105±4µmol quanta $m^{-2}$ $s^{-1}$; medium:333±16 µmol quanta $m^{-2}$ $s^{-1}$ and high: 657±11 µmol quanta $m^{-2}$ $s^{-1}$. **Temperature**: 23ºC; **salinity**: 35‰ | 21 days | branchlets of 10 cm length | >6 replicates | Mexican Caribbean | *S. fluitans* | 0.028 to 0.11 | 32 |
| | 13 light levels ranging from 8–576 | **Two experiment replicates. Light levels:** from 8 to 576 µE $m^{-2}$ $s^{-1}$**.** **Temperature:** no specific; **salinity** 36‰ | 21 days | 20 mm of length | 6 replicates (3 in each experiment) | Neritic region (Florida Waters) | *S. natans* (I) | 0.0 to 0.0727 ± 0.0049 | 31 |
| | Three light treatments: using a 12:12 h light: dark photoperiod. | **Two experiment replicates:** August-September of (1) 2019 and (2) 2020. **Light levels:** Low:105±4µmol quanta $m^{-2}$ $s^{-1}$; medium:333±16 µmol quanta $m^{-2}$ $s^{-1}$ and high: 657±11 µmol quanta $m^{-2}$ $s^{-1}$. **Temperature**: 23ºC; **salinity**: 35‰ | 21 days | branchlets of 10 cm length | >6 replicates | Mexican Caribbean | *S. natans* (I) | 0.023 to 0.051 | 32 |
| Response to temperature | Four temperature treatments: 12, 18, 24, 30 ºC | **Light conditions:** 110 µE $m^{-2}$ $s^{-1}$ with 14:10 h light–dark. **Temperature range:** 12, 18, 24, 30 ºC**;** **Salinity:** 36‰ | 21 days | 20 mm of length | 6 replicates | Neritic region (Florida Waters) | *S. fluitans* | 0 to 0.048 ± 0.02 | 31 |
| | Two temperature treatments: ~28 and ~31ºC | Two experiments replicates: July and September 2019 **Light conditions:** 435 and 581 µmol $m^{-2}$ $s^{-1}$; temperature range: ~28 and ~31; **Salinity:** 35‰. | 20 days | 12 g wet weight | 4 replicates | Mexican Caribbean | *S. fluitans* | 0.04 ± 0.01 to 0.06 ± 0.001 | 33 |
| | Four treatments: 22, 25, 28, 31 ºC | Three experimental series: (1) April-may, (2) July and (3) October to November 2021. **Light conditions**: between 435 and 581 µmol quanta $m^{-2}$ $s^{-1}$; **Temperature range**: 22, 25, 28, 31 ºC; **salinity**: 35‰ | (1) 43 days, (2) 18 days, and (3) 30 days | No specific | 4-7 replicates | Mexican Caribbean | *S. fluitans* | 0.058 ± 0.01 to 0.095 ± 0.01 | 34 |
| | Two treatments: ~27.6±0.4 and ~29.6±0.3 | **Two periods:** May and August-September 2022. **Light conditions:** Natural irradiances; **temperature:** ~27.6±0.4 and ~29.6±0.3; **salinity:** 32±2‰ | 7 days in each period | ~40 g wet weight | 33 replicates (15, 18) | Tropical Atlantic | *S. fluitans* | 0.06 ± 0.02 to 0.098 ± 0.01 | 35 |
| | Four treatments: 21,23,27, 31ºC | **Light conditions:** 210±57 µmol $m^{-2}$ $s^{-1}$ using 12:12 h light:dark cycle.**Temperature range**: 21,23,27, 31ºC ; **salinity range:** 36‰. | 5 days | No specific | 10 replicates | Neritic region | *S. fluitans* | 0.049 ± 0.01 to 0.073 ± 0.01 | 36 |
| | Four treatments: 22, 25, 28, 31 ºC | **Three experimental series:** (1) April-may, (2) July and (3) October to November 2021. | 1) 43 days, (2) 18 days, | No specific | 4-7 replicates | Mexican Caribbean | *S. natans* (I) | 0.054 ± 0.01 to 0.067 ± 0.01 | 34 |

| | | | | | | | | | |
|---|---|---|---|---|---|---|---|---|---|
| | | **Light conditions:** between 435 and 581 µmol quanta $m^{-2}$ $s^{-1}$ ; **temperature range**: 22, 25, 28, 31 ºC; **salinity**: 35‰. | and (3) 30 days | | | | | | |
| | Two treatments: ~27.6±0.4 ºC and ~29.6±0.3ºC | Two periods: May and August-September 2022. **Light conditions:** Natural irradiances; **temperature:** ~27.6±0.4 and ~29.6±0.3; **salinity:** 32±2‰ | 7 days in each period | ~40 g wet weight | 33 replicates (15, 18) | Tropical Atlantic | *S. natans* (I) | 0.042 ± 0.02 to 0.051 ± 0.01 | 35 |
| | Four treatments: 21,23,27, 31ºC | **Light conditions: 210**±57 µmol $m^{-2}$ $s^{-1}$ using 12:12 h light:dark cycle.**Temperature range**: 21,23,27, 31ºC ; s**alinity range:** 36‰. | 5 days | No specific | 10 replicates | Neritic region | *S. natans* (I) | 0.046 ± 0.01 to 0.072 ± 0.01 | 36 |
| | Four treatments: 22, 25, 28, 31 ºC | Three experimental series: (1) April-may, (2) July and (3) October to november 2021. **Light conditions: Bewteen 435 and 581 µmol quanta $m^{-2}$ $s^{-1}$ ; temperature range**: 22, 25, 28, 31 ºC; **salinity**: 35‰. | (1) 43 days, (2) 18 days, and (3) 30 days | No specific | 4-7 replicates | Mexican Caribbean | *S. natans* (VIII) | 0.045 ± 0.01 to 0.059 ± 0.01 | 34 |
| | Two treatments: ~27.6±0.4 and ~29.6±0.3 | **Two periods**: May and August-September 2022. **Light conditions:** Natural irradiances; **temperature:** ~27.6±0.4 and ~29.6±0.3; **salinity:** 32±2‰ | 7 days in each period | ~40 g wet weight | 33 replicates (15, 18) | Tropical Atlantic | *S. natans* (VIII) | 0.029 ± 0.02 to 0.035 ± 0.01 | 35 |
| | Two temperature treatments: ~28 and ~31ºC | **Two experiments replicates:** July and September 2019. **Light conditions:** 435 and 581 µmol $m^{-2}$ $s^{-1}$; **temperature range:** ~28 and ~31; s**alinity:** 35‰. | 20 days | 12 g wet weight | 4 replicates | Mexican Caribbean | *S. natans* (VIII) | 0.05 ± 0.02 to 0.055 ± 0.01 | 33 |
| | Four treatments: 21,23,27, 31ºC | **Light conditions: 210**±57 µmol m-2 s-1 using 12:12 h light:dark cycle.**Temperature range**: 21,23,27, 31ºC ; s**alinity range:** 36‰. | 5 days | No specific | 10 replicates | Neritic region | *S. natans* (VIII) | 0.014 ± 0.01 to 0.040 ± 0.01 | 36 |
| Response to fertilization (ex situ) | Added seawater enriched and control | Added seawater enriched in $NO_3^-$ ; $NH_4^+$; $PO_4^{3-}$ and control (not enrichment). **Light conditions:** 39–60 µE $m^{-2}$ $d^{-1}$ ; **temperature**: 28-30ºC; **salinity**: 36‰ | 7-10 days | 50 g wet weight | 2 replicates for each treatment | Oceanic region | *S. fluitans* | 0.03 ± 0.006 to 0.056 ± 0.007 | 37 |
| | Added seawater enriched and control | Added seawater enriched in $NO_3^-$ ; $NH_4^+$; $PO_4^{3-}$ and control (not enrichment). **Light conditions:** 48–60 µE $m^{-2}$ $d^{-1}$ ; **temperature**: 28-30ºC; **salinity**: 36‰ | 7-10 days | 50 g wet weight | 2 replicates for each treatment | Neritic region | *S. fluitans* | 0.036 ± 0.007 to 0.061 ± 0.007 | 37 |
| | Addition of fertilizer in high and low concentrations and control. | **Two experiments replicates**: Feb-March 2017 and April - May 2017. Addition of fertilizer with 6.6% $NH_4^+$, 8.4% $NO_3^-$, 9% P2O5, and 12% K in high and low concentrations and control (not enrichment). **Light conditions:** attenuation of <60% of natural irradiance; **temperature**: 27-29ºC; **salinity**: 36‰ | 21 days | 40–50 g wet weight | 2–6 replicates for each treatment | Mexican Caribbean | *S. fluitans* | 0.012 ± 0.004 to 0.039 ± 0.002 | 33 |
| | Added seawater enriched and control | Added seawater enriched in $NO_3^-$ ; $NH_4^+$; $PO_4^{3-}$ and control (not enrichment). **Light conditions: 39**–60 µE $m^{-2}$ $d^{-1}$ ; **temperature**: 28–30ºC; **salinity**: 36‰. | 7-10 days | 50 g wet weight | 2 replicates for each treatment | Oceanic region | *S. natans* (I) | 0.029 ± 0.005 to 0.055 ± 0.004 | 37 |

| | | | | | | | | | |
|---|---|---|---|---|---|---|---|---|---|
| | Added seawater enriched and control | Added seawater enriched in $NO_3^-$ ; $NH_4^+$; $PO_4^{3-}$ and control (not enrichment). **Light conditions:** 48–60 µE $m^{-2}$ $d^{-1}$ ; **temperature**: 28–30ºC; **salinity**: 36‰. | 7-10 days | 50 g wet weight | 2 replicates for each treatment | Neritic region | *S. natans* (I) | 0.043 ± 0.009 to 0.072 ± 0.008 | 37 |
| | Addition of fertilizer in high and low concentrations and control. | **Two experiments replicates:** Feb-March 2017 and April - May 2017. Addition of fertilizer with 6.6% $NH_4^+$, 8.4% $NO_3^-$, 9% $P_2O_5$, and 12% K in high and low concentrations and control (not enrichment). **Light conditions:** attenuation of <60% of natural irradiance; **temperature**: 27-29ºC; **salinity**: 36‰ | 21 days | 40–50 g wet weight | 2–6 replicates | Mexican Caribbean | *S. natans* (VIII) | 0.038 ± 0.004 to 0.056 ± 0.002 | 33 |
| | | **Two experiments replicates:** May 2022. Four different nutrient treatments. Seawater enriched with 100 µM of $NO_3^-$ + 10 µM of $PO_4^{3-}$; 10 µM of Fe ; both (N,P + Fe) and control (not enrichment). **Light conditions:** 327 to 824 µmol $m^2$ $s^{-1}$ ; **temperature:** 28±1ºC; **salinity:** 35‰ | 5.5 to 7.3 days | 5.5–6 g wet weight | 4 replicates for each treatment in each experiment | Mexican Caribbean | *S. fluitans* | 0.059 to 0.184 | 38 |
| Response to salinity (ex situ) | Salinity range: 12‰, 18‰, 24‰, 30‰, 36‰, 42‰ | **Two experiments repeated. Light conditions**: 110 µE $m^{-2}$ $s^{-1}$ with 14:10 h light–dark; **temperature**: no specific; **salinity range:** 12‰, 18‰, 24‰, 30‰, 36‰, 42‰. | 21 days | 20 mm of length | 6 (3 replicates in each treatment) | Neritic region (Florida Waters) | *S. fluitans* | Data not shown | 31 |
| | Salinity range: 28‰, 32‰, 36‰, 40‰ | **Light conditions: 210**±57 µmol $m^{-2}$ $s^{-1}$ using 12:12 h light:dark cycle.**Temperature**: 25-26ºC during day and 27.5 to 29.9ºC during night; s**alinity range:** 26‰, 28‰, 30‰, 36‰, 38‰, 40‰. | 5 days | No specific | 10 replicates for each treatment | Neritic region (Florida Waters) | *S. fluitans* | 0.027 ± 0.01 to 0.054 ± 0.01 | 36 |
| | Salinity range: 12‰, 18‰, 24‰, 30‰, 36‰, 42‰ | Two experiments repeated. Light conditions: 110 µE $m^{-2}$ $s^{-1}$ with 14:10 h light–dark; **temperature**: 25-26 during day and 27.5 to 29.9 during night; s**alinity range:** 12‰, 18‰, 24‰, 30‰, 36‰, 42‰. | 21 days | 20 mm of length | 6 (3 replicates in each treatment) | Neritic region (Florida Waters) | *S. natans* (I) | 0 to 0.049 ± 0.01 | 31 |
| | Salinity range: 28‰, 32‰, 36‰, 40‰ | **Light conditions:** 210±57 µmol $m^{-2}$ $s^{-1}$ using 12:12 h light:dark cycle; **temperature**: 25-26ºC during day and 27.5 to 29.9ºC during night; **salinity range:** 26‰, 28‰, 30‰, 36‰, 38‰, 40‰. | 5 days | No specific | 10 replicates for each treatment | Neritic region (Florida Waters) | *S. natans* (I) | 0.010 ± 0.01 to 0.054 ± 0.01 | 36 |
| | Salinity range: 28‰, 32‰, 36‰, 40‰ | **Light conditions: 210**±57 µmol $m^{-2}$ $s^{-1}$ using 12:12 h light:dark cycle; **temperature**: 25-26ºC during day and 27.5 to 29.9ºC during night; **salinity range:** 26‰, 28‰, 30‰, 36‰, 38‰, 40‰. | 5 days | No specific | 10 replicates for each treatment | Neritic region (Florida Waters) | *S. natans* (VIII) | 0.037 ± 0.01 to 0.041 ± 0.01 | 36 |
| Characteri-zation of regional populations | Natural conditions | Sampling between March 1986 and June 1990. **Light conditions:** Natural irradiances conditions; temperature: no specific; salinity: 36‰. | 4-6 days | 20 g wet weight | 6 replicates | Neritic caribbean | *S. fluitans* | 0.04 ± 0.01 to 0.05 ± 0.01 | 39 |
| | Natural conditions | Sampling between March 1986 and June 1990. **Light conditions:** Natural irradiances conditions; temperature: no specific; salinity: 36‰. | 4-6 days | 20 g wet weight | 6 replicates | Neritic N. Atlantic | *S. fluitans* | 0.03 ± 0.01 to 0.093 ± 0.01 | 39 |

| | | | | | | | | |
|---|---|---|---|---|---|---|---|---|
| Natural conditions | **Sampled between may-june 2021.** **Light conditions:** 74 to 740 µmol photons $m^{-2}$ $s^{-1}$; **temperature**: 28-31ºC. | 9 days | 20 g wet weight | 12 replicates | Eastern Caribbean | *S. fluitans* | 0.019 ± 0.01 to 0.063 ± 0.02 | 40 |
| Natural conditions | Sampling between March 1986 and June 1990. **Light conditions:** Natural irradiances conditions; temperature: no specific; salinity: 36‰. | 4-6 days | 20 g wet weight | 6 replicates | Neritic caribbean | *S. natans (I)* | 0.004 ± 0.001 | 39 |
| Natural conditions | Sampling between March 1986 and June 1990. **Light conditions:** Natural irradiances conditions; temperature: no specific; salinity: 36‰. | 4-6 days | 20 g wet weight | 6 replicates | Neritic N. Atlantic | *S. natans (I)* | 0.05 ± 0.005 to 0.091 ± 0.01 | 39 |
| Natural conditions | Sampling between March 1986 and June 1990. **Light conditions:** Natural irradiances conditions; temperature: no specific; salinity: 36‰. | 4-6 days | 20 g wet weight | 6 replicates | Oceanic Sargasso Sea | *S. natans (I)* | 0.01 ± 0.005 to 0.02 ± 0.005 | 39 |
| Natural conditions | **Sampled between may-june 2021.** **Light conditions:** 74 to 740 µmol photons $m^{-2}$ $s^{-1}$; **temperature**: 28-31ºC. | 9 days | 20 g wet weight | 12 replicates | Eastern Caribbean | *S. natans* (I) | 0.018 ± 0.02 to 0.029 ± 0.02 | 40 |
| Natural conditions | **Sampled between may-june 2021.** **Light conditions:** 74 to 740 µmol photons $m^{-2}$ $s^{-1}$; **temperature**: 28-31ºC. | 9 days | 20 g wet weight | 12 replicates | Eastern Caribbean | *S. natans* (VIII) | 0.008 ± 0.01 to 0.043 ± 0.01 | 40 |

**Table S3.** Literature data for the variation of thallus size (blade area) of the three holopelagic *Sargassum* variants present in the GASB. Units for algal size are expressed in $cm^2$ $thallus^{-1}$. Mean±Es and the number of replicates are indicated in parenthesis. N.D = no data.

| Specie | Oceanic region | Location | Collection date | Specimens' status | Thallus size ($cm^2$ $thallus^{-1}$) | N | Source Ref. |
|---|---|---|---|---|---|---|---|
| *S. natans* (I) | Caribbean Sea | Puerto Morelos, Mexico | 2018 | Wild | 0.38 ± 0.01 | 50 | This study |
| | | South coast of Costa Rica | 2019 | Wild | 0.19 ± 0.8 | 10 | 41 |
| | | Humacao, Puerto Rico | 2021 | Wild | 0.30 ± 0.01 | 15 | 42 |
| *S. natans* (I) | Western Tropical Atlantic | Oceanic stations | 2014 | Wild | 0.60 ± 0.07 | 15 | 8 |
| | | Oceanic stations | 2015-2016 | Wild | 0.29 ± 0.02 | 8 | 7 |
| *S. natans* (I) | West Africa | Ondo State, Nigeria | 2019-2020 | Wild | 0.26 ± 0.0 | N.D. | 43 |
| *S. natans* (VIII) | Caribbean Sea | Puerto Morelos, Mexico | 2018 | Wild | 1.63 ± 0.08 | 50 | This study |
| *S. natans* (VIII) | Western Tropical Atlantic | Oceanic stations | 2014 | wild | 2.35 ± 0.1 | 85 | 8 |
| *S. fluitans* (III) | Caribbean Sea | Cienfuegos, Cuba | may-2012 | Wild | 0.26 ± 0.02 | 10 | 12 |
| | | Puerto Morelos, Mexico | 2018 | Wild | 0.68 ± 0.02 | 50 | This study |
| | | Puerto Morelos, Mexico | 2019-2020 | Wild | 0.73 ± 0.01 | 44 | 32 |
| | | Puerto Morelos, Mexico | 2019-2020 | Laboratory | 0.80 ± 0.04 | 40 | 32 |
| | | Punta Yabuca, Puerto Rico | 09-dic-21 | Wild | 0.36 ± 0.03 | 12 | 44 |
| *S. fluitans* (III) | Western Tropical Atlantic | Oceanic stations | 2014 | wild | 0.99 ± 0.01 | 85 | 8 |
| | | Oceanic stations | 2015-2016 | wild | 0.90 ± 0.09 | 10 | 7 |
| *S. fluitans* (III) | West Africa | Ondo, State, Nigeria | 2019-2020 | Wild | 0.72 ± 0.0 | N.D | 43 |

**Table S4.** Linear and non-linear (power-functions) for the associations of variation between optical and structural traits of holopelagic *Sargassum*. Bold highlights significant correlations.

| Y, X, trait | Specie | $R^2$ | Ecuation | *F*-statistic | Df | $P_{value}$ |
|---|---|---|---|---|---|---|
| $A_{PAR}$ vs. Total [Chl*a* + Chl*c*] content | *S. natans* (I) | 0.22 | Y = 46.81±5.83 + 0.29±0.16x | 3.26 | 11 | 0.098 |
| | *S. natans* (VIII) | 0.02 | Y = 46.15±6.54 + 0.08±0.20x | 0.160 | 7 | 0.700 |
| | *S, fluitans* (III) | 0.03 | Y = 55.22±5.80 – 0.10±0.15x | 0.438 | 14 | 0.518 |
| $A_{Chla}$ vs. Chl*a* content | ***S. natans* (I)** | **0.48** | **Y = 32.06±6.08 + 0.75±0.22x** | **11.49** | **12** | **0.005** |
| | *S. natans* (VIII) | 0.14 | Y = 30.03±13.37 + 0.63±0.57x | 1.188 | 7 | 0.312 |
| | *S, fluitans* (III) | 0.02 | Y = 51.65±8.63 + 0.14±0.28x | 0.224 | 14 | 0.643 |
| $a^*_{PAR}$ vs. Total [Chl*a* + Chl*c*] content | ***S. natans* (I)** | **0.74** | **$Y = 0.42 \pm 0.19 * X^{(-0.80 \pm 14)}$** | **26.9** | **11** | **0.003** |
| | ***S. natans* (VIII)** | **0.80** | **$Y = 0.57 \pm 0.26 * X^{(-0.96 \pm 22)}$** | **25.44** | **7** | **0.001** |
| | ***S. fluitans* (III)** | **0.78** | **$Y = 1.53 \pm 0.96 * X^{(-1.21 \pm 0.18)}$** | **42.62** | **14** | **0.001** |
| $a^*_{Chla}$ vs. Chl *a* content | ***S. natans* (I)** | **0.59** | **$Y = 0.297 \pm 0.14 * X^{(-0.72 \pm 0.15)}$** | **2.712** | **12** | **0.018** |
| | *S. natans* (VIII) | 0.22 | $Y = 0.263 \pm 0.41 * X^{(-0.74 \pm 0.52)}$ | 1.902 | 7 | 0.210 |
| | *S, fluitans* (III) | 0.47 | $Y = 0.60 \pm 0.52 * X^{(-0.90 \pm 0.26)}$ | 0.202 | 14 | 0.66 |

**Table S5**. ANCOVA tests for investigating significant differences in the associations of variation between optical traits and pigment content per projected within each variant. Models are based on log-log transformations of the optical and pigment content direct values described in plots C and D of figure 3, which illustrate: (C) the non-linear association between the specific absorption coefficient for the average PAR range (400-700 nm; $a^*_{pigm}$), and total chlorophyll Chl[*a*+*c*] density (content pero projected area); and (D) the non-linear association between the specific absorption coefficient of chlorophyll *a* at 675 nm, $a^*_{Chla}$, and Chl*a* density (content per projected area).

| Source | Df | SM | MS | F-ratio | $P_{value}$ |
|---|---|---|---|---|---|
| Log Total chlorophyll content | 1 | 2.4955 | 2.4955 | 97.8847 | 2.943e-11* |
| Factor. Species | 2 | 0.3142 | 0.1571 | 6.1619 | 0.005** |
| Log Total chlorophyll content*Species | 2 | 0.0806 | 0.0403 | 1.5810 | 0.22 |
| Residuals | 32 | 0.8158 | 0.0255 | | |
| **Source** | **Df** | **SM** | **MS** | **F-ratio** | $P_{value}$ |
| Log Chl *a* content | 1 | 1.0580 | 1.0580 | 19.78 | 9.281e-0.5* |
| Factor. Species | 2 | 0.3093 | 0.1546 | 2.8924 | 0.0695 |
| Log Chl *a* content*Species | 2 | 0.0664 | 0.0332 | 0.6213 | 0.5434 |
| Residuals | 33 | 1.7645 | 0.05347 | | |

**Table S6**. Analysis of variance (ANOVA tests) for investigating differences in the relative growth rates (RGR) values documented in the literature, for the three holopelagic *Sargassum* variants present in the Central tropical Atlantic.
Sf (III) = *Sargassum fluitans* (III); Sn (I) = *Sargassum natans* (I); Sn (VIII) = *Sargassum natans* (VIII).

| Source | DF | SS | MS | *F*-statistic | *P*-value |
|---|---|---|---|---|---|
| Total | 191 | 0.2113 | 0.0011 | | |
| A | 2 | 0.0486 | 0.0243 | 28.22 | < 0.001 |
| Error | 189 | 0.1627 | 0.00086 | | |
| **Tukey's all pairs comparison** | | | | | |
| **Comparison** | | **Mean differences** | ***T*-value** | ***P*-value** | **95% confidence** |
| Sf (III) vs. Sn (VIII) | | 0.03566 | 8.7108 | < 0.001 | 0.022 to 0.0493 |
| Sf (III) vs. Sn (I) | | 0.0297 | 8.9027 | < 0.001 | 0.018 to 0.0409 |
| Sn (I) vs. Sn (VIII) | | 0.0058 | 1.3983 | 0.5849 | -0.008 to 0.012 |

**Sources**: Hanisak & Samuel[31], Magaña-Gallegos et al[34,] Magaña-Gallegos et al[33,34], Corbin & Oxenford[35], Schell et al[36], Lapointe[37], Leemans et al[38], Lapointe et al[39], Changeux et al[40]
